# Design and self-assembly of cytomotive filaments

**DOI:** 10.64898/2026.01.07.698144

**Authors:** Marija Krstić, Christian Vanhille-Campos, Buzz Baum, Maitane Muñoz-Basagoiti, Anđela Šarić

## Abstract

Cytomotive filaments such as actin and FtsZ exhibit treadmilling, a turnover mode in which subunits are added at one end and removed from the other, driven by nucleotide hydrolysis. This dynamics, which requires directional filament growth and disassembly without fragmentation, is crucial for the function of these filaments. However, how treadmilling is encoded in the monomer properties remains unclear. Here we combine physical modelling with *in silico* evolution to identify the minimal monomer design principles that yield treadmilling. We show that directional growth requires a polymerization-induced change in the binding interface of the monomer. Fragmentation-free disassembly further requires that this conformational change allosterically couples two monomer interfaces that bind it to its left and right neighbours. Guided by these principles, we design a minimal particle-based monomer that spontaneously assembles into treadmilling filaments in simulations. This work establishes a physical framework for cytomotive filament dynamics and provides a foundation for engineering synthetic systems with controllable emerging treadmilling dynamics.

**Significance statement:** Treadmilling, the continuous, energy-driven assembly and disassembly of monomers at opposite ends of a filament, is a fundamental dynamic of the cytoskeleton across the tree of life. How can individual monomers self-assemble into a polar filament that grows at one end and shrinks at the other? By combining physical modelling and in silico evolution, we uncover the physical and geometrical principles that monomers must satisfy to naturally form treadmilling filaments. In particular, we show that polar assembly requires monomers to undergo a conformational change upon assembly. In turn, controlled filament disassembly requires this conformational change to geometrically couple a monomer’s two binding interfaces, preventing filament fragmentation. Our work provides a physical framework for designing functional self-assembling materials.

## I. INTRODUCTION

Cytomotive filaments, which comprise the actin and tubulin superfamilies, are characterised by polarity and turnover: they preferentially grow in one direction and, powered by nucleotide hydrolysis, cycle between assembly and disassembly [1, 2]. As a result, they are able to perform functional work in the cell [3, 4]. The function of filaments critically depends on their specific polymerisation dynamics [5–8]. For example, by transitioning between growth and shrinkage periods, microtubules assist chromosome separation during mitosis [7, 9, 10]. Likewise, bacterial cell division and forces driving eukaryotic cell motility rely, respectively, on FtsZ and actin treadmilling [11–13], a specific polymerisation dynamics whereby filaments predominantly grow on one end and shrink on the other.

It has long been postulated that treadmilling dynamics arises from the combination of polar self-assembly at equilibrium, and the turnover of monomers driven by nucleotide hydrolysis [14]. However, connecting the structural and chemical properties of single monomers to their emergent cytomotive polymerisation and depolymerisation kinetics remains an open challenge. Crystallographic data shows that cytomotive proteins like actin and FtsZ adopt two different conformations in the polymer and in solution [1, 15, 16]. These conformations have been hypothesized to contribute to avoiding filament fragmentation [1]. Experimental and simulation results further indicate that cytomotive proteins exploit nucleotide hydrolysis (ATP for actin, and GTP for FtsZ) to destabilize binding interfaces, and drive monomer turnover [17–20]. Nonetheless, how the monomer structure translates into asymmetric polymerisation kinetics, and contributes to directional depolymerisation of the filament is still unclear.

In this work, we identify the physical and geometrical principles of monomers that self-assemble into treadmilling filaments by combining minimal physical models, computer simulations and *in silico* evolution. Our key result is that polar filament growth and polar filament shrinking, the defining features of treadmilling dynamics, require monomers to switch conformation upon assembly through a deformation that allosterically couples the two ends of a monomer that bind it to its neighbours. By identifying essential features of cytomotive self-assembly, our work establishes a physical framework to design emergent functional kinetics from the monomer properties.

## II. RESULTS

We start by introducing the previous literature-established requirements for treadmilling dynamics [14]. Polar growth implies faster growth on the filament head compared to the tail (Fig. 1A). In single-stranded filaments monomer binding is thermodynamically equivalent at both filament ends, hence polar growth must stem from different energy barriers for monomer binding on the filament head versus tail. Furthermore, for a monomer to shift its thermodynamic preference from assembling to disassembling, energy must be dissipated, for example, through nucleotide hydrolysis. Hydrolysis is a stochastic process whose probability increases with monomer residence time within the polymer, thereby naturally establishing a gradient of hydrolysed monomers toward the slower-growing end. If the filament predominantly disassembles at its ends (Fig. 1B), and does not fragment in the middle, the energy-driven cycling of monomers between assembly and disassembly results in treadmilling (Fig. 1B, C). Since these dynamics emerge from monomer interactions alone, treadmilling must necessarily be encoded in the physical and geometrical properties of the single monomers themselves, which we investigate in what follows.

**FIG. 1.**
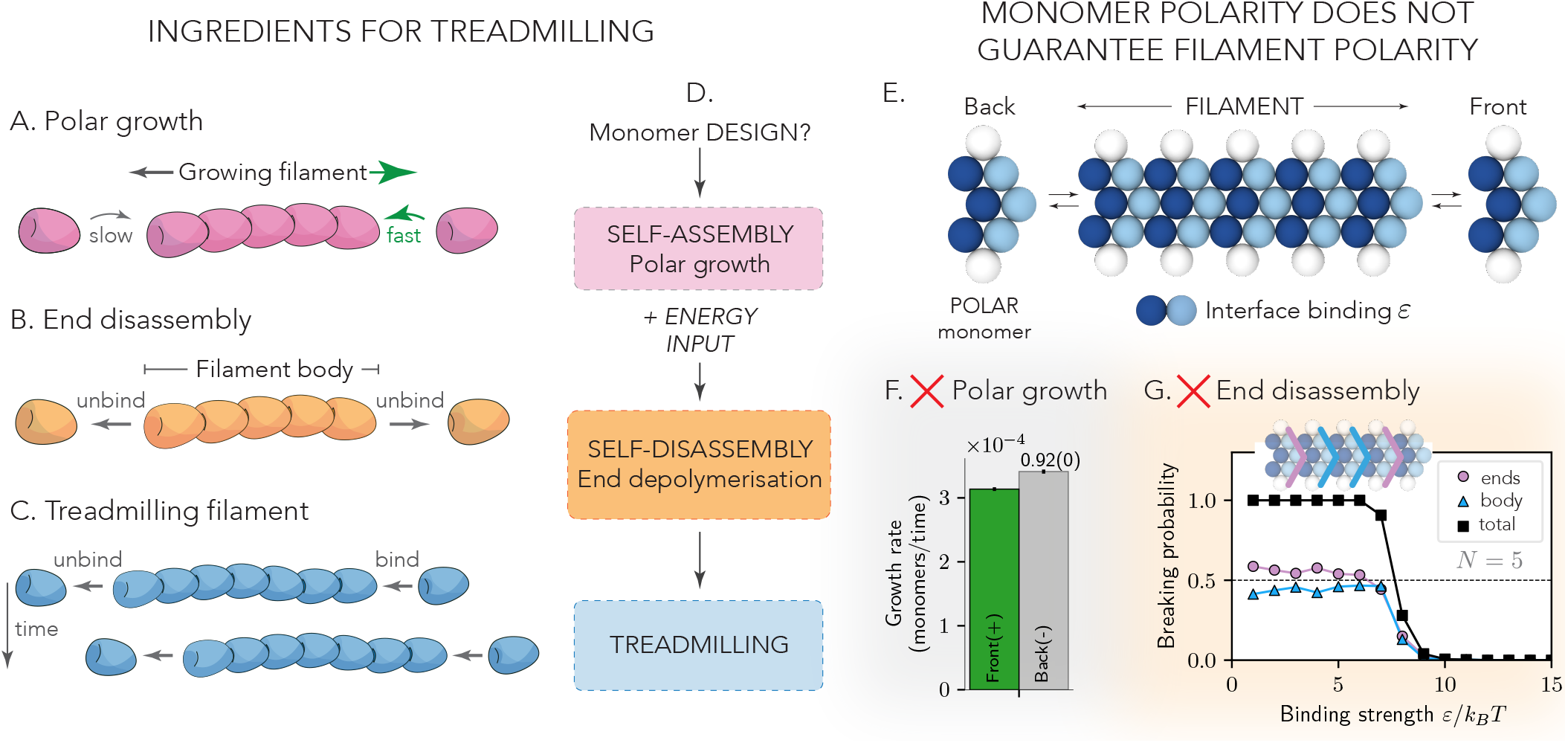
From monomer design to polymerisation dynamics. Treadmilling dynamics naturally emerge when polar assembly is combined with end disassembly by energy dissipation. A) Filament exhibiting polar growth, characterised by faster addition on one end than the other. B) Filament exhibiting end disassembly: monomers dissociate from the ends without fragmentation. C) Emergent treadmilling: a filament grows on one end while disassembling on the other, without fragmenting. D)Schematic illustrating how the ingredients of treadmilling arise from the monomer design in combination with energy influx.E) Example system of a polar colloidal monomer forming filaments. F) Polar growth is not recovered by such a system: the growth rate is the same on both ends (*r*_+_*/r* _*−*_ = 0.92(0), *n*_replica_ = 300). G) End disassembly is not allowed in such a system: filaments exhibit the same probability of fragmenting and end-shrinking when interface strength is reduced from the stable non-shrinking regime. The black curve shows percentage of simulations where disassembly occurs: the filaments are stable for *ε >* 9*k*_*B*_*T* (*n*_replica_ = 300). See Supplementary Information for simulation details.

### A. structurally polar monomer does not give rise to treadmilling

We first consider the assembly of structurally polar monomers with distinguishable front and back interfaces. We build such monomers out of attractive and volume-excluded beads connected by springs (Fig. 1E), which interact with binding with energy *ε* to self-assemble into linear polymers. See Supplementary Information (SI) for details.

To determine the potential of our monomers to form treadmilling filaments, we design two Molecular Dynamics (MD) simulation protocols aimed at evaluating filament assembly and disassembly dynamics: the polar growth test, and the fragmentation test. In the polar growth test, we simulate the self-assembly of polymers from a dimer seed at constant monomer concentration to probe for asymmetric binding rates. In the filament fragmentation test, we record the location of the first unbinding event in a pre-assembled filament to quantify the probability of filament fragmentation as the binding strength between monomers decreases.

The MD simulations reveal that our structurally polar monomers readily self-assemble into filaments, which however show no polar growth (Fig. 1F). Furthermore, by comparing unbinding probabilities of a monomer in the body of the filament versus those at the filament end (Fig. 1G), we find that no energetic regime enables preferential end-depolymerisation compared to filament fragmentation for this monomer design. This demonstrates that monomer polarity does not guarantee filament polarity, nor does it enable controlled disassembly.

### B. Evolving filament polarity

To gain insight into which monomer properties might yield polar growth we start with an agnostic monomer structure, and use Genetic Algorithm (GA) [21–23] to evolve monomers that exhibit polar self-assembly kinetics.

We consider a genotypic space spanned by 4*×* 4 particles in the hexagonal lattice connected by stiff bonds (Fig. 2A) and use a loss function that evaluates polymerisation asymmetry to select for monomers that yield filaments with kinetic polarity. To promote linear assemblies, we place the interface-forming binding sites at opposite sides of the lattice, and allow sites of different types to interact with each other through a short-range attractive potential, while the rest of the particles interact via volume exclusion. We evaluate the fitness of each design using MD simulations (see SI for further details). Our GA results, illustrated by two example designs in Fig. 2B (TAIL and CAPPER), show that evolved monomers can successfully self-assemble into polar filaments with non-negligible kinetic asymmetry (see Suppl.Movies 1 and 2).

**FIG. 2.**
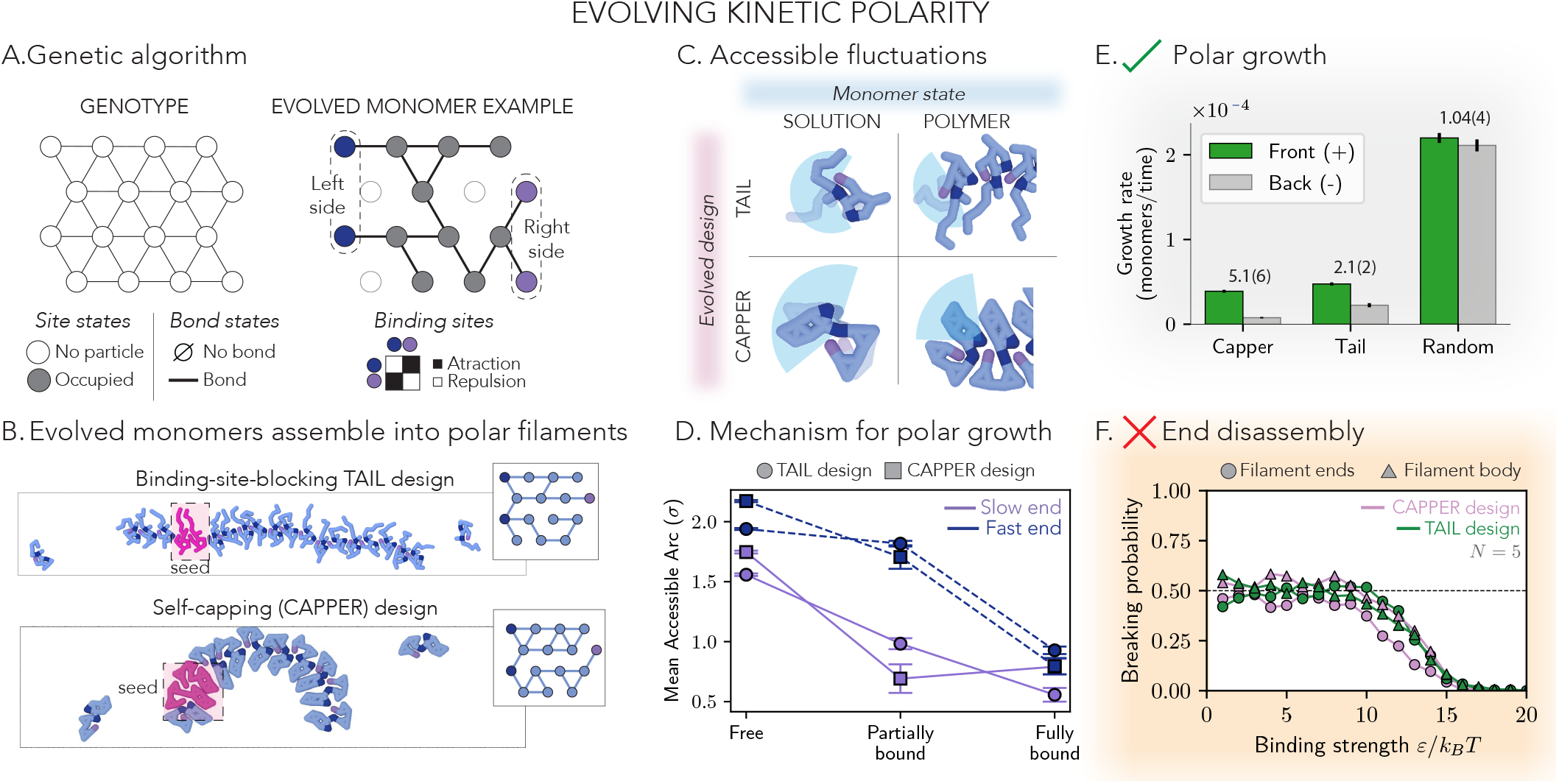
Evolving monomer designs that exhibit polar growth upon assembly. A) Left: Illustration of the monomer genotype, with particle sites that can be empty or occupied and interparticle bonds that can be present or not. Right: Example of an evolved monomer design with binding sites in blue and purple defining the interfaces and gray particles for the core of the monomer. B) Two examples of evolved monomer designs and the emergent self-assembled polymers they form. Filaments exhibit polar growth: more monomers are added to the right of the seed (pink) than the left at a given time *t* (Suppl. Movies 1 and 2). C) The monomer fluctuations are affected by its polymer binding. Accessible fluctuations of the monomer structure are highlighted by blue arcs, which decrease as monomers are incorporated to a polymer. D) Polar growth is a direct result of emergent asymmetries in the accessible binding area between the two monomer interfaces upon binding. When monomers are partially bound (filament edge), a difference in mean accessible arc between the two sides emerges. E) Evolved monomer designs exhibit polar growth rates (faster on one side than the other, i.e., *r*_+_ *> r*_*−*_ ), while random monomer designs do not, *r*_+_ *∼ r*_*−*_ (*n*_replica_ = 300). F) Evolved monomer designs do not exhibit end disassembly: no binding affinity enables a preference for filament ends depolymerisation over filament fragmentation (*n*_replica_ = 300). See SI for further simulation details.

We find that configurational fluctuations are a hallmark of successful evolved designs that can be exploited for polar self-assembly. Monomers exhibit substantial configurational fluctuations in solution. Polymerisation restricts these fluctuations and results in a reduced accessible binding area when a monomer is bound in the filament (Fig. 2C). In a sense, the *average configuration* of the monomer is different in solution or in a polymer.Crucially, we find that this accessible binding area reduction strongly depends on the side of the monomer that binds to the filaments, such that one filament side will display slower binding kinetics (Fig. 2D). Specifically, the slow side is the one where the edge monomer has the least accessible surface area. Looking more closely into the TAIL design in the polymer and solution states (Fig 2C), we find that the fluctuating monomer arm (or tail) gets localised around a binding site in the polymer state, inducing a decrease in the available binding area and therefore affecting further polymerization on that end. Similarly, for the CAPPER design, polymerization imposes restricted fluctuations of the monomer conformation such that one of the binding sites is buried, preventing further polymerisation on that end. Critically, these effects are localised to only one end of the filament. Indeed, our MD simulations show that evolved designs exhibit polar filament growth, while random monomer designs before the evolution do not (Fig. 2E). Interestingly, we find that these monomer designs do not en-able controlled end-disassembly (Fig. 2F), thus failing to spontaneously produce treadmilling filaments when coupled to stochastic binding affinity changes.

Together, these results demonstrate that polar self-assembly requires monomers that fluctuate between multiple conformational states and in particular adopt distinct configurations upon polymerization compared to solution, which matches experimental observations in actin and FtsZ [1, 15, 16]. However, these results also show that a polymerization-associated conformational switch is not sufficient to enable treadmilling, as it fails to localise disassembly to the filament ends, contrary to what has been previously hypothesized [1]. In what follows we investigate what additional constraints need to be met by multistate monomers in order to achieve both polar assembly and controlled end-disassembly, and thus exhibit emergent treadmilling.

### C. Monomers with a conformational switch build polar filaments

To systematically investigate the role of monomer conformational switch in treadmilling, we next develop the simplest possible monomer model that can exist in multiple conformations, namely, a monomer that fluctuates along a single coordinate around only two well-defined states: a solution and a polymer state. This minimal model allows for both simulation and analytical tractability, unlike the complex fluctuating structures evolved in the previous section.

To build our minimal monomer, we consider three rigid bars of length *x*_0_ and four binding sites: a horizontal *base* with a binding site at each end, a vertical *support* fixed at a distance *x* along the base, and a *hinging arm* that can rotate around the support, with a binding site at each end (Fig. 3A; Suppl. Movie 3). Two breakable torsional springs control the configurational fluctuations of the monomer, with two well-defined stable states: a solution (*S*) form and a polymer (*P* ) form. We define the *P* form such that it can self-assemble into linear filaments (Fig. 3B) while the *S* form will correspond to a higher opening angle *θ* of the hinging arm. Binding sites interact via an attractive potential (Fig. 3B), which will modify the energy of the monomer in the polymer compared to solution (Fig. 3C).

**FIG. 3.**
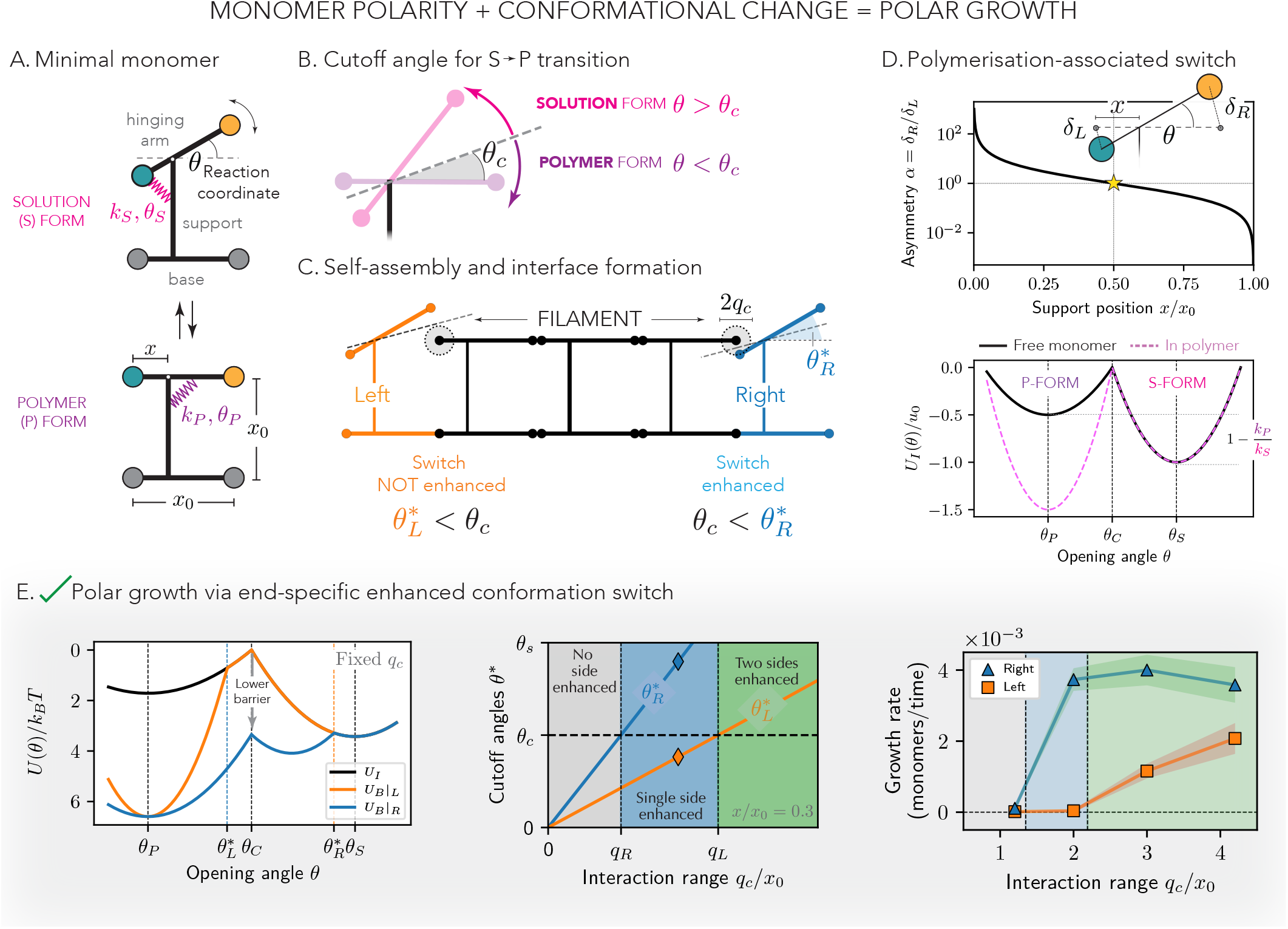
What kind of conformational change enables treadmilling? Developing a minimal model exhibiting polar growth. A) Minimal monomer model consisting of three rigid bars connecting four binding sites. The hinging arm (top bar) can rotate around the support (vertical bar), fixing the geometry via the angle *θ* (Suppl. Movie 3). Fluctuations between a solution form (S-form, pink) and a polymer form (P-form, green, parallel bars) are controlled by breakable torsional springs with constants *k*_*S*_ and *k*_*P*_, and equilibrium rest angle *θ*_*S*_ and *θ*_*P*_, respectively. B) The cutoff angle *θ*_*c*_ dictates the state of the monomer. C) Monomers can only form polymers in the P-form. If the S-form is polar (asymmetric), the binding sites interaction will be felt at different points along the *S→ P* transition depending on the binding side (left, orange, vs right, blue). 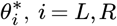, *i* = *L, R* sets the opening angle at which the left and right monomer, respectively, enter within interaction range *q*_*c*_ of the polymer. In the schematic, the monomer from the right (blue) interacts with the polymer already at the *S* state, i.e., 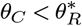, while the monomer from the left (orange) must transition from *S* to *P* in order to bind the polymer, i.e., 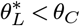 . D) The support position determines the asymmetry of the polymerisation-associated switch and the polarity of the S-form (top).The binding sites attraction can stabilise the P-form, promoting polymerisation (bottom). E) S-form polarity can result in end-specific catalysis of the binding reaction for the right interaction range regime, decreasing the activation barrier for the S-form to P-form transition and leading to faster self-assembly kinetics on one end of the filament (Suppl. Movie 4). This is confirmed in simulations (right panel, *n*_replica_ = 200 per point; shaded area gives the standard error of the mean).

The hinging arm rigidly couples the two interfaces of the monomer, such that a displacement of the left binding site induces a displacement of the right one, and viceversa. The position of the support *x* dictates the amplitudes of the interface deformations at each side (*δ*_*L*_,*δ*_*R*_) as the monomer switches between *S* and *P* conformations. The amplitude of the deformations is equal only for symmetric monomers (*x* = 1*/*2, Fig. 3C).

A natural consequence of this amplitude asymmetry in polar monomers is that the opening angle *θ*^*\**^ for which binding sites come within interaction range *q*_*c*_ of the polymer will be different on either side of the monomer (Fig. 3B). The energy landscape of an incoming monomer can be hence very different at polymer head compared to the polymer tail (Fig. 3D). As a result, the polymer can act as an effective catalyser for monomer binding on one side only, decreasing the activation barrier for the *S→ P* reaction.This will naturally result in different binding kinetics at the polymer head versus polymer tail.

For asymmetric monomers (*x* ≠ 1*/*2) then, three kinetic regimes emerge depending on the monomer geometry (*x* and *θ*_*S*_) and interaction range (Fig. 3D): (*i*) binding is not catalysed on either side of the monomer; (*ii*) binding is catalysed only for the monomer binding from its short hinging arm side; (*iii*) binding is catalysed on both sides of the monomer. In regime (*ii*) we thus expect asymmetric monomers to self-assemble into polar filaments, with faster growth rates at the front (where free monomers bind from their short hinging arm side) than the back. Indeed, this picture is confirmed in MD simulations of asymmetric toy monomers for different interaction ranges, where we clearly observe the three catalysis regimes (Fig. 3D). Together, these results demonstrate that a structurally polar (asymmetric) conformational change in monomers between a solution and polymer configurations enables polar self-assembly of filaments, in agreement with our GA results. Furthermore, our minimal approach reveals how effective, end-specific, binding catalysis can be encoded in monomer structure and exploited to tune self-assembly kinetics.

Next we turn to filament stability and disassembly. If structural monomers that switch between configurations can exhibit kinetic polarity, what enables stable filament disassembly without fragmentation, once binding affinity changes? What was missing from our evolved designs?

### D. Allosteric two-state monomers disfavour fragmentation

To localise monomer disassembly to the filament ends, and prevent filament fragmentation, the properties of a monomer must change depending on its position within the filament. At the assembly level, the key distinction between a monomer within the filament, and a monomer at its ends is the number of binding neighbours — the former has two, while the latter has one. Based on this simple geometrical rule, we next study how the difference in the number of binding interfaces can be leveraged to promote the stability of the filament body and localise disassembly to the filaments ends with monomers that switch between configurations.

Building on our previous results, we now examine the stability of bound minimal monomers positioned either at the filament end or within the filament core. Hydrolysis-driven reductions in binding affinity can make the polymer form less stable than the solution form, thereby promoting disassembly (Fig 4A). In what follows, we show how this stability transition can manifest differently when a monomer is bound on only one side, and how this asymmetry can be leveraged to facilitate end-localized disassembly.

**FIG. 4.**
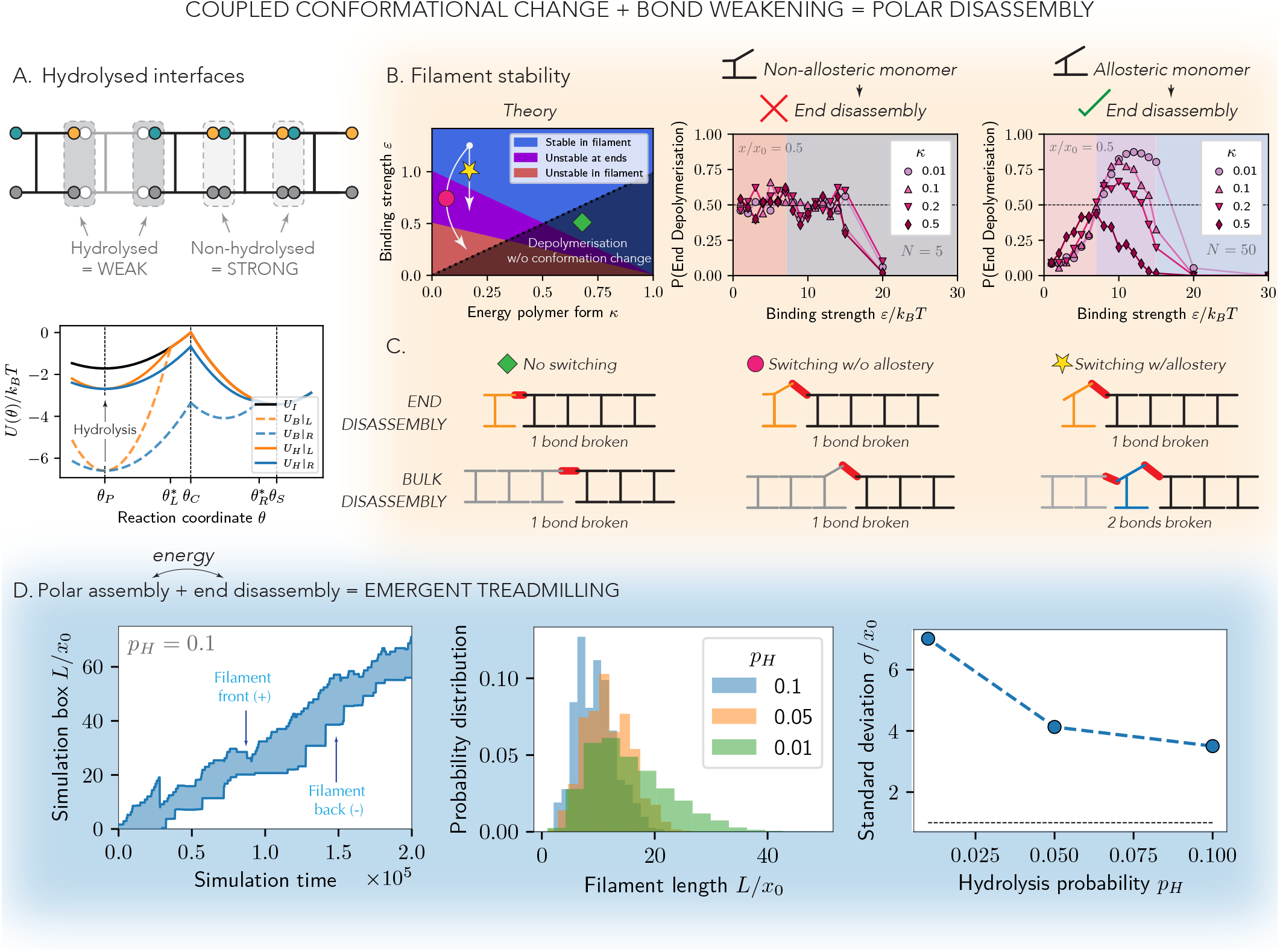
Polar disassembly requires allosteric monomers with a conformational switch. A) Hydrolysis decreases the affinity of binding sites, destabilising monomer-monomer interfaces. B) Two-state monomers with coupled interfaces exhibit end-specific disassembly in the right binding strength regime. Simulations for non-allosteric and allosteric monomers confirm that coupled interfaces are a necessary condition for end disassembly (*n*_replica_ = 50 per parameter set). The parameter *κ* = *k*_*P*_ */k*_*S*_ controls the stability of the *P* -form. C) Schematics of the mechanism enabling end disassembly in allosteric twostate monomers. Fragmentation requires breaking two interfaces, while end-disassembly only requires breaking one. If the interfaces are decoupled or the monomer does not change conformation this effect is lost. D) Treadmilling naturally emerges in simulations of minimal two-state allosteric and polar monomers when stochastically hydrolysing interfaces at a given rate *p*_*H*_ . Left: Example trajectory of a treadmilling filament (Suppl. Movie 4). Middle: Filament length distributions for different hydrolysis rates. Right: Filament length standard deviation as a function of the hydrolysis rate (*n*_replica_ = 50 per hydrolysis rate). See SI for further simulation details.

We show both in theory and simulations that in a certain range of *ε* binding strength values of hydrolysed interfaces we can promote end disassembly, while keeping monomers in the filament body stable (purple region in Fig 4 B). Critically, we show that this regime only emerges when monomers exhibit a polymerization-associated switch and this switch allosterically couples the two monomer interfaces (Fig 4 B-C).

If monomers exhibit no polymerization-associated switch, which we enforce in our minimal model by constraining the monomer to its polymer form only, unbinding requires breaking a single interface regardless of the monomer’s position within the filament. The kinetics of fragmentation and end-disassembly are thus equivalent for all binding affinities, such that no end-localised disassembly is possible (Fig 4 C, left). Conversely, if monomers exhibit a polymerization-associated switch, they can unbind by switching to the solution state. This transition requires breaking two interfaces in the filament body but only one at the filament end (Fig 4 C, right). This implies the polymer state is more stable in the filament body compared to the ends for a given binding affinity (Fig 4 A, bottom). We show in simulations that this enables the existence of an optimal binding affinity that maximises end-disassembly compared to fragmentation (Fig 4 B, right). Finally, we show that this mechanism requires allostery between the two binding interfaces (Fig 4 B-C, middle). If we allow binding sites to fluctuate independently of each other (floppy hinging arm), we find no binding affinity regime where end-disassembly is favoured, as unbinding in the filament body can now happen by breaking a single interface. In light of these results, we postulate that the complex GA evolved designs are likely failing to treadmill because they lack this rigid interface allostery.

Given that our minimal two-state model (Fig 3A) exhibits both kinetic polarity and filament stability, we next examine its treadmilling capacity, coupling filament growth with nucleotide hydrolysis. Informed by the stability analysis, we choose the hydrolysed binding strength to be in the region where only end monomers of the filament are destabilised. We start from short filaments with constant monomer concentrations on the two ends and a fixed hydrolysis rate. This gives rise to emergent tread-milling dynamics, where we can control the distribution of filament lengths by tuning the hydrolysis probability (Fig 4D; Suppl. Movie 5).

Altogether, the results of this section show that preventing filament fragmentation after nucleotide hydrolysis (i.e., a reduction in binding affinity) requires a conformational switch that allosterically couples the two monomer interfaces. Combined with the earlier conclusion that an asymmetric, polymerization-associated conformational switch is necessary for polar self-assembly (Section C), we can now identify the essential requirements for a two-state monomer to treadmill: it must switch between a structurally polar solution state and a polymer state while allosterically coupling its two interfaces. Importantly, our simulations demonstrate that these conditions are sufficient for treadmilling to emerge spontaneously.

### E. Emergent treadmilling in a colloidal monomer design

Let us now return to the initial structurally polar colloidal monomer design (Fig. 1E) and demonstrate how one can, incorporating the design principles uncovered in the previous sections, design colloidal monomers with controllable emergent treadmilling dynamics.

We start by removing one of the internal bonds of the monomer, such that it can now fluctuate between two distinct configurations: a triangle-like *solution* form, that cannot form polymers, and the original chevron-like *polymer* form (Fig 5 A). These fluctuations are controlled by the binding affinity of two orthogonal pairs of particles (Fig 5 A), akin to our minimal model from previous sections. We show that this monomer design displays considerable growth rate asymmetry in our standard kinetic polarity test (Fig 5 B; Suppl. Movie 6). This asymmetry arises because the available binding area is larger on one side of the solution state than the other, such that a free polymer end is found first on that side, resulting in faster binding kinetics (Fig 5 C). In summary, with the conformational switch we introduce a solution form with asymmetric binding areas, leading to kinetic polarity.

**FIG. 5.**
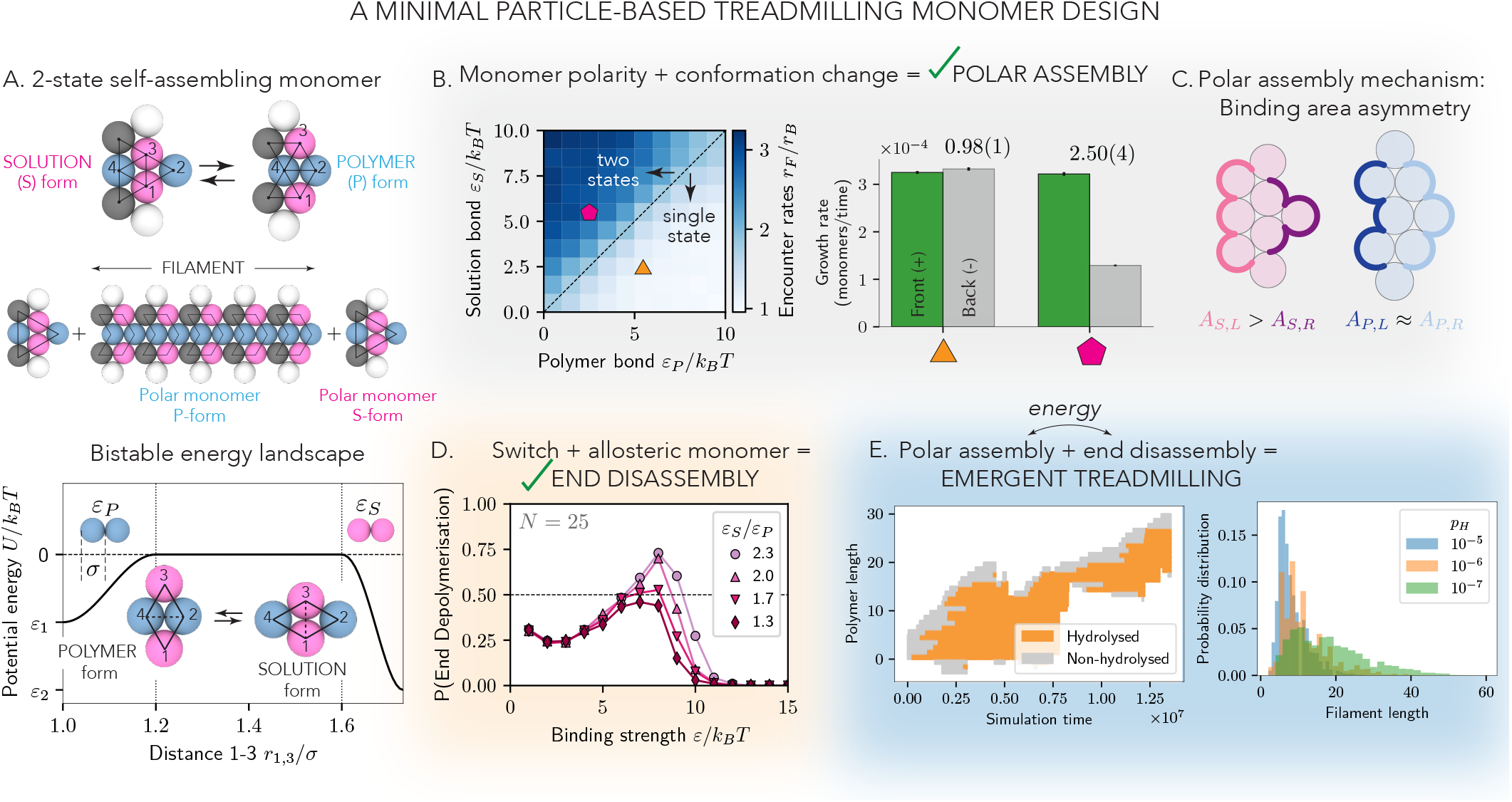
Particle-based treadmilling monomer design. A) Two-state self-assembling monomer design. Monomers fluctuate between two well-defined conformations, controlled by the affinity of the pink and blue particle pairs respectively. Only one conformation (P-form) can form polymers. B) Polar assembly spontaneously emerges for monomers that are preferentially in the S-form when unbound (stronger pink affinity). C) Polar growth is mediated by an asymmetry in the exposed binding area of the S-form between the tip (purple) and back (pink). This results in asymmetric assembly kinetics driven by polar encounter rates. D) Because the two monomer interfaces are coupled (monomer allostery), a binding strength regime exists where end disassembly without fragmentation is favoured (*n*_replica_ = 300). E) Treadmilling naturally emerges in simulations when stochastically hydrolysing interfaces to the end disassembly regime at a given rate *p*_*H*_ . Left: Example trajectory of a treadmilling filament. Right: Filament length distributions for different hydrolysis rates.

Next, we consider the disassembly kinetics of the new design. The conformational switch we describe allosterically couples the two interfaces, satisfying the conditions described in the previous section, and should thus enable end-only disassembly for some binding affinity regime. Indeed, filament stability tests show a region of binding strengths where end disassembly is favoured (Fig 5 D).

Finally, we demonstrate that this design exhibits emergent self-assembly treadmilling dynamics in simulations where we stochastically weaken interfaces to model nucleotide hydrolysis (Fig 5E; Suppl. Movie 7). Again, the hydrolysis binding strength is set to the binding strength that maximizes end disassembly for a given internal landscape. Altogether, this analysis demonstrates that the design principles identified from our minimal monomer model translate directly to more complex colloidal structures. Specifically, if a monomer can induce binding area asymmetry by switching between a polymer state and solution state, its two interfaces are allosterically coupled, and stochastic binding-affinity reduction occurs within the polymer, then treadmilling will emerge – regardless of the monomer’s specific nature. We illustrate this with a simple colloidal monomer design, showing that these principles are general and can be readily exploited to engineer monomers that self-assemble into treadmilling structures with tunable properties.

## III. DISCUSSION

In this work, we investigate the structural features that must be encoded at the monomer level to enable self-assembly into treadmilling filaments. For a filament to exhibit treadmilling dynamics, it needs to support polar growth while maintaining structural integrity under hydrolysis, i.e., without undergoing fragmentation. These two criteria serve as the basis for evaluating the tread-milling potential of monomer designs explored in this study.

Using a combination of methods – genetic algorithms, an analytically tractable minimal model, MD simulations and complex colloidal monomer designs – we have identified the essential conditions under which simple monomers robustly self-assemble into treadmilling filaments. Rather than depending on the detailed chemistry or architecture of the building block, sustained tread-milling requires a set of well defined physical ingredients, which are both necessary and sufficient for its spontaneous emergence (Fig 6):

**FIG. 6.**
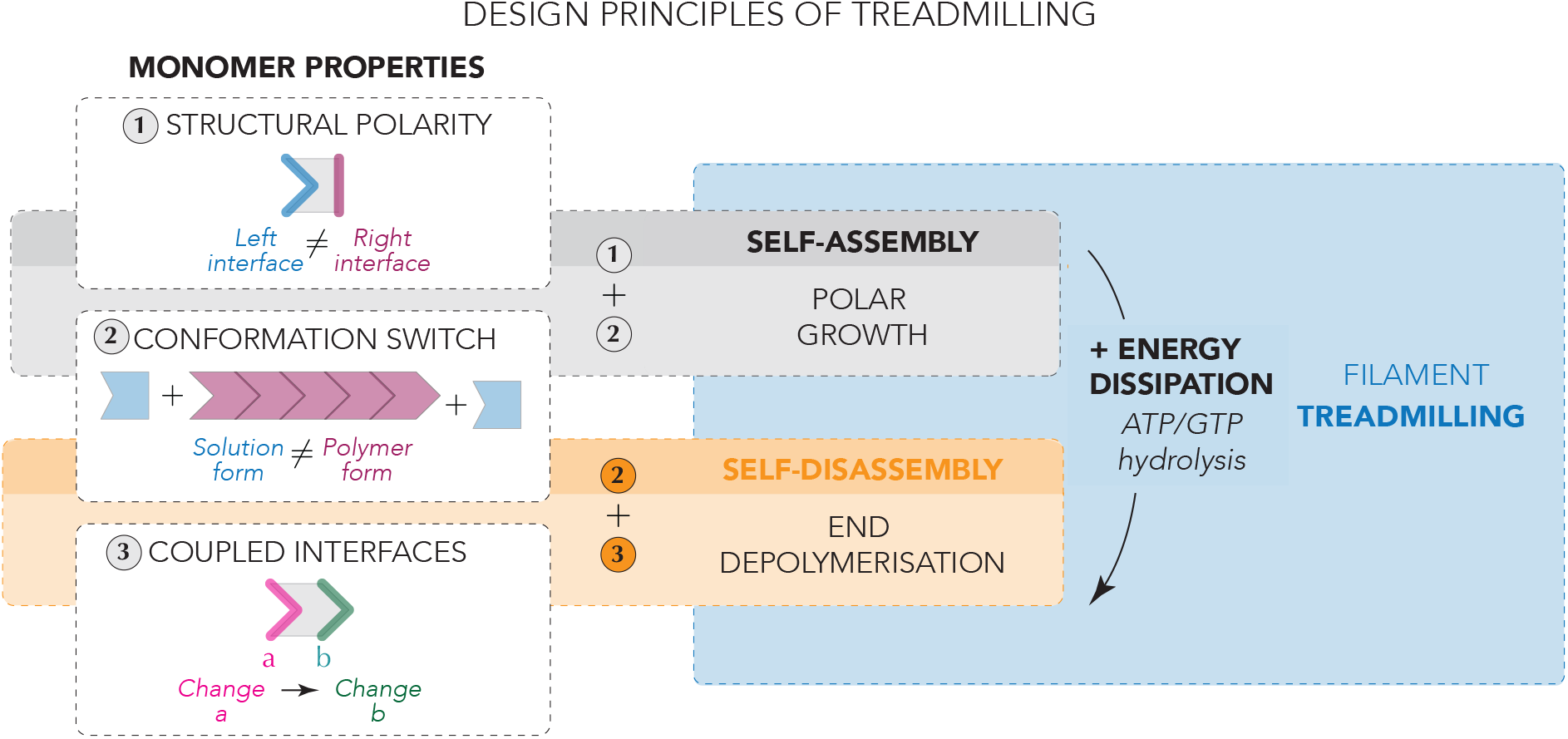
Design principles of treadmilling. There are three key monomer properties that, in combination, give rise to treadmilling dynamics. On one hand, a conformational switch between a solution form and a polymer form combined with structural polarity of the solution form produces polar growth at equilibrium (1 + 2, gray). On the other hand, if the solution-polymer conformational switch couples the two monomer interfaces (i.e., a switch on one side results in a switch on the other), a binding strength regime emerges where disassembly can be localised to the filament ends, avoiding fragmentation (2 + 3, orange). Coupling these two regimes via energy influx by means of nucleotide hydrolysis, which affects the interface binding affinity, spontaneously produces treadmilling filaments that grow on one end and shrink on the other without fragmenting (1+ 2 + 3 + energy dissipation, blue).

1. The monomer must induce binding area asymmetry on the polymer ends by switching between a polymer state and solution state;
2. Its two binding interfaces must be allosterically coupled; and
3. A stochastic reduction of binding affinity must occur within the polymer (e.g., through hydrolysis).

Together, these features ensure polarized assembly and controlled, end-localized disassembly without filament fragmentation. Importantly, we demonstrate the generality of these principles by constructing a simple colloidal monomer that implements them. Molecular dynamics simulations confirm that this design self-assembles into treadmilling filaments with tunable dynamics, showing that the principles uncovered here extend directly to more complex architectures.

This work underscores the central role of a polymerization-associated conformational switch in cytomotive filament dynamics. Based on experimental observations, a conformation switch had been previously hypothesized as the mechanism to prevent filament fragmentation upon nucleotide hydrolysis [1, 16]. Here we confirm this hypothesis by explicitly showing that this switch is essential both for directional assembly as well as ensuring filament stability during disassembly.

The emergence of directional assembly via a switch through monomer fluctuations in our evolved designs is notable in light of the intrinsically disordered regions identified in treadmilling FtsZ near both the C- and N-termini of the protein [24, 25]. Our findings suggest that these regions may play a direct role in regulating filament kinetics through steric interactions alone.

The design principles identified in this work provide a framework for engineering synthetic cytoskeletal filaments in synthetic cell systems, illustrating how macro-molecular structure and interactions can be tuned to generate complex collective behaviour. By directly linking monomer architecture to polymer polarity, our results also clarify the minimal requirements for directional dynamics in cytomotive systems. Unlike multistranded filaments, where sequential kinetics and lateral interactions can contribute to emergent directionality, single-stranded filaments cannot readily exploit such collective effects [1], highlighting the necessity of the principles developed here. Future work could explore how these design rules extend to more complex cytoskeletal assemblies and whether they can be used to recapitulate diverse forms of polymer turnover and dynamics

Our modelling framework is readily translatable to experimental systems based on programmable interactions, enabling direct implementation of these principles. In particular, our designs can be realized using platforms such as DNA-coated colloidal particles [26–28], synthetic peptides, or RNA-based technologies. These approaches open the door to the creation of self-assembling, directional polymers at the microscale and establish a foundation for engineering synthetic cytomotility in biomimetic and artificial systems.

## ACKNOWLEDGEMENTS

We thank Ferdinand Horvath for assisting in the rendering of protein structures. CVC acknowledges funding by the European Union’s Horizon 2020 Research and Innovation Programme under Marie Skłodowska-Curie Grant Agreement No. 101278760. MM-B acknowledges funding by the European Union’s Horizon 2020 research and innovation programme under Marie Skłodowska-Curie Grant Agreement No. 101034413. MK acknowledges funding by the Gesellschaft für Forschungsförderung Niederösterreich (GFF) under the RTI-Dissertations 2024. This work was supported by the Royal Society (Grant No. UF160266; CVC and AŠ) and the European Union’s Horizon 2020 Research and Innovation Programme (Grant No. 802960; CVC and AŠ).

