## Supplementary Information for "Design and self-assembly of cytomotive filaments"

### Design and self-assembly of cytomotive filaments
#### *Supplementary Information*

Marija Krstić[1,*], Christian Vanhille-Campos[1,2,3,*],
Buzz Baum[4], Maitane Muñoz-Basagoiti[1], Anđela Šarić[1]

[1]Institute of Science and Technology Austria, Klosterneuburg, Austria
[2]Department of Physics and Astronomy, Institute for the Physics of Living Systems, University College London, London, United Kingdom
[3]Laboratoire Jean Perrin, CNRS/Sorbonne Université, Paris, France
[4]MRC Laboratory of Molecular Biology, Cambridge, UK
[*]M.K. and C.V.C contributed equally to this work. First authorship order is alphabetic.

June 19, 2026

**4 Implementation of the genetic algorithm** **20**

**5 Supplementary Movies** **25**

### 1 Elastic network model

In this section, we describe the building block design introduced in Figs. 3 and 4 in the Main Text. The model is designed to be minimal yet analytically tractable, and therefore uses an elastic network representation based on breakable harmonic springs.

#### 1.1 Defining the polymer subunit

We consider a two-dimensional building block made of three bars of length $\sigma = 1$ connecting four binding sites at the corners of a square, defined as follows:

- A rigid horizontal bar (*base*) connecting binding sites at points $(0,0)$ and $(1,0)$.
- A rigid vertical bar (*support*) at a distance $x$ along the *base*, with $x \in (0,1)$.
- A rigid bar (*hinging arm*) connecting the binding sites at (0, 1) and (1, 1).

The *hinging arm* can rotate around the top of the *support*, and consequently, change the position of the top binding sites. The rotation is controlled by torsional springs (see subsection 1.1.1). As a result, the geometric state of the monomer is defined by the angle $\theta$, namely, the angle that the *hinging arm* forms with the horizontal, which we use as the reaction coordinate. See Fig. 3A in Main Text for a schematic of the building block.

##### 1.1.1 The torsional harmonic springs and reaction coordinate

We introduce two torsional harmonic springs to construct a double-well energy landscape along the reaction coordinate $\theta$ (Fig. 3C in the Main Text). Each spring is characterized by a well-defined equilibrium angle, which sets the orientation of the hinging arm and consequently, the specific geometric state of the monomer. In particular,

- The closed or polymer ($P$) state is characterised by
  - Equilibrium angle $\theta_P = 0$ (or equivalently, $\theta_P = \pi/2$, depending on the reference point).
  - Harmonic oscillation around the equilibrium angle

$$U(\theta) = \frac{1}{2}k_P(\theta - \theta_P)^2 \text{ if } \theta < \theta_C \text{ and 0 elsewhere}$$

- The open or solution ($S$) state is characterised by
  - Equilibrium angle $\theta_S > \theta_P$
  - Harmonic oscillations around the equilibrium angle

$$U(\theta) = \frac{1}{2}k_S(\theta - \theta_S)^2 \text{ if } \theta > \theta_C \text{ and 0 elsewhere}$$

We choose the cutoff angle $\theta_C$ such that when the solution spring breaks, the polymer spring forms, or conversely, such that when the polymer spring breaks, the solution spring forms. This way, the two harmonic potentials along $\theta$ do not overlap. We also choose the $P$-form geometry as the one that the monomer adopts when it is within the polymer.

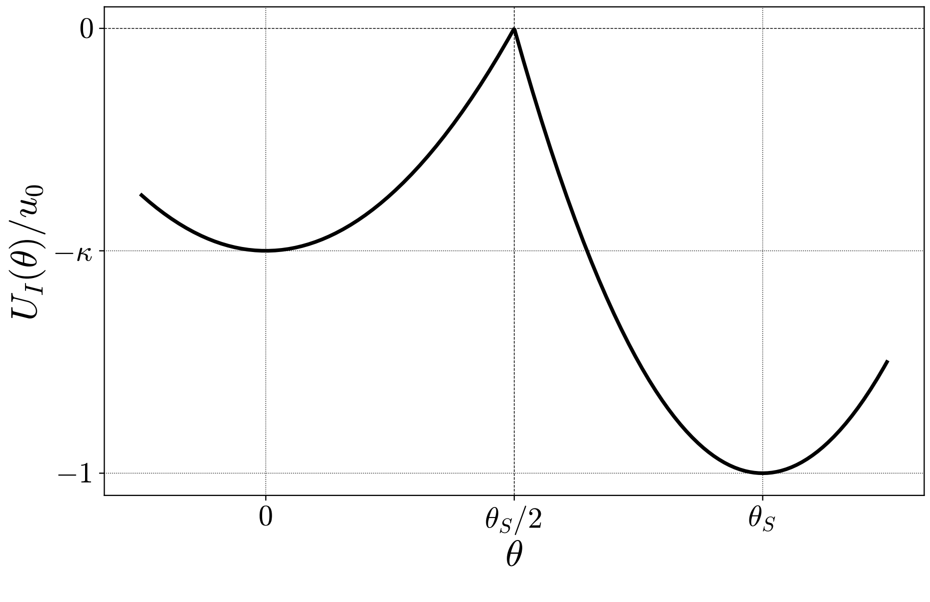

Supplementary Figure 1: Example internal energy landscape $U_I(\theta)$ for a monomer in our elastic network model. Here $\kappa = 0.5$, $k = 50k_BT$, and $\theta_S = \pi/3$ yielding $u_0 = 6.85\ k_BT$.

###### 1.1.2 The internal energy landscape of the building block

The internal energy landscape of the building block results from combining the two torsional springs above, and imposing the continuity of the potential at $\theta_C$ (see Supp. Fig. 1). This results in

$$U_I(\theta) = \frac{k}{2}\left[H\left(\frac{\theta_S}{2} - \theta\right)\kappa\left(\theta^2 - \frac{\theta_S^2}{4}\right) + H\left(\theta - \frac{\theta_S}{2}\right)\left((\theta - \theta_S)^2 - \frac{\theta_S^2}{4}\right)\right], \tag{1}$$

where $H$ is the Heaviside function, $k = k_S$ is the spring constant of the solution spring and $\kappa = k_P/k_S$ is the ratio between the polymer and solution spring constants. For simplicity we set the cutoff angle $\theta_C = \theta_S/2$, where the potential vanishes. Altogether, the double-well harmonic potential is fully defined by only three variables, namely, $k$, $\kappa$ and $\theta_S$. For convenience, the potential in Eq. 1 is also shifted, i.e., $U(\theta_S/2) = 0$, such that its value at the two minima $\theta_S$ and $\theta_P$ coincides with the activation barriers to switch between monomer states. The value of the minima correspond to $U(\theta_S) = -u_0$ and $U(\theta_P) = -\kappa u_0$, where $u_0 = \frac{1}{8}k\theta_S^2$ is the characteristic energy scale of the system. The energy difference between the polymer and solution states is

$$\Delta U_I = U_I^P - U_I^S = (1 - \kappa)u_0. \tag{2}$$

The solution state is more stable than the polymer state when $\kappa < 1$, which is necessary for kinetic polarity. If $\kappa > 1$, the monomer does not have to change conformation upon polymerisation, making it impossible to establish asymmetric binding kinetics at either side of the filament, as the left and right monomer interfaces would be identical in the absence of any interactions already.

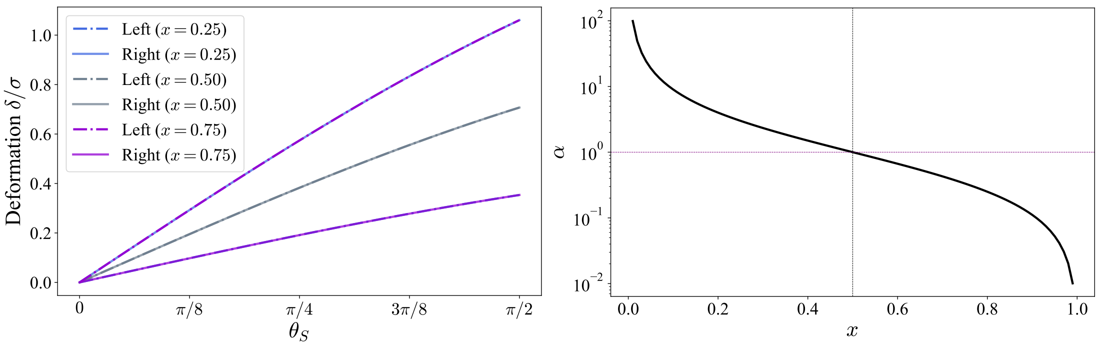

Supplementary Figure 2: (Left) Interface deformations on either side as a function of $\theta_S$ for different values of $x$. Only when $x = 0.5$ are the deformations on both sides equal. (Right) Monomer asymmetry $\alpha$ as a function of the support position $x$.

##### 1.1.3 Asymmetry of the building block

The position of the top binding sites within the monomer depends on the angle $\theta$. We define $\delta$ as the distance that each binding site must traverse for the monomer to go from its solution state ($\theta = \theta_S$) to its polymer state ($\theta = \theta_P$). Depending on whether the binding sites are located on the left (L) or right (R) of the monomer, we have

$$\delta_L^2 = 2(1 - \cos\theta_S)x^2 \qquad \text{and} \qquad \delta_R^2 = 2(1-\cos\theta_S)(1-x)^2. \tag{3}$$

We define the asymmetry of the monomer as the ratio between the right and left deformations, which is independent of the opening angle, as

$$\alpha = \frac{\delta_R}{\delta_L} = \frac{1-x}{x}. \tag{4}$$

For $x = 0$ or $x = 1$ the interfaces become decoupled, and $\alpha$ becomes ill-defined, as only one of the two interfaces deforms upon monomer conformation switch. Note also that $\alpha$ is symmetric around $x = 1/2$ upon inversion of the two sides, meaning that $\alpha(x) = 1/\alpha(1-x)$ or that $\delta_L(x) = \delta_R(1-x)$ for any choice of $\theta_S$ (see Fig. 3C in the Main Text and Supp. Fig. 2).

#### 1.2 Defining the formation of interfaces

We now describe the self-assembly of the building blocks introduced in section 1.1. For that, let us consider an attractive potential between binding site pairs of equal color. For simplicity, we use again a harmonic potential:

$$U_B(q) = \frac{1}{2}k_I\left(q^2 - q_c^2\right) \text{ if } q < q_c \text{ and 0 elsewhere} \tag{5}$$

Here $k_I$ defines the binding strength of the potential, $q$ is the inter-site distance, which is a function of $x$ and $\theta$ as we show below, and $q_c$ is the interaction range. The minimum of the potential corresponds to

$$U_B(0) = -\frac{1}{2}k_I q_c^2, \tag{6}$$

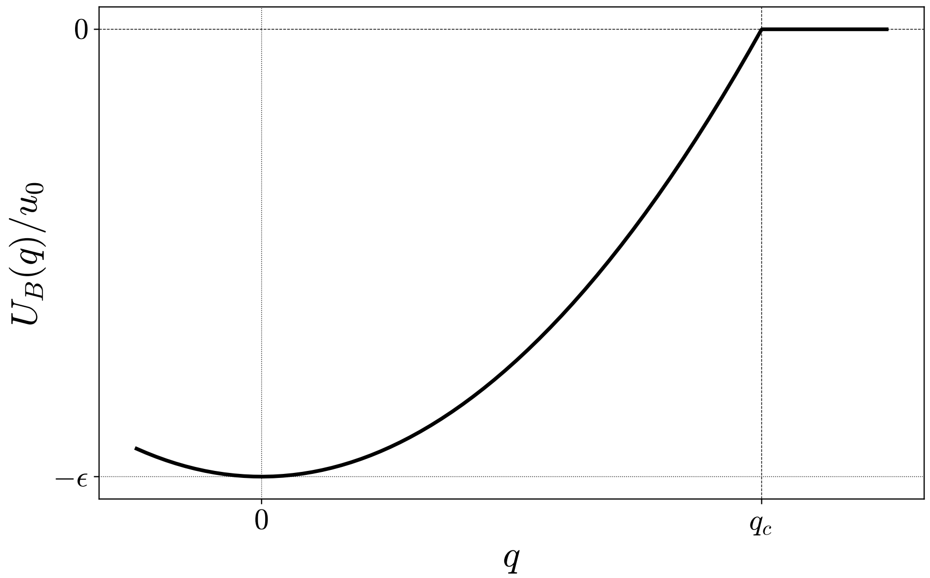

Supplementary Figure 3: Example binding potential $U_B(q)$ for $\epsilon = 2$ and $q_c = 0.25\sigma$. Here $k = 50k_BT$ and $\theta_S = \pi/3$ such that $k_I = 685.38k_BT/\sigma^2$.

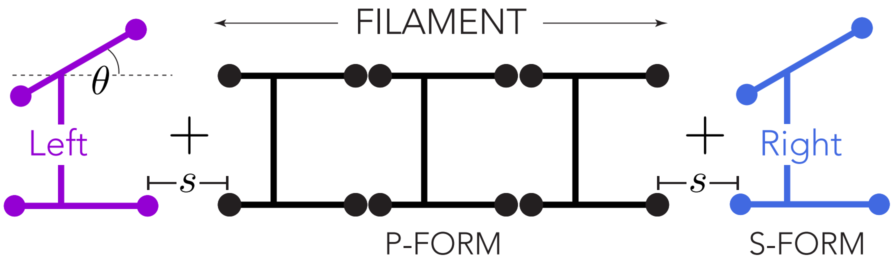

Supplementary Figure 4: Schematic of the binding of two monomers to an existing polymer, illustrating the two relevant degrees of freedom: inter-monomer distance $s$ and opening angle $\theta$.

which can be rewritten in terms of the energy scale $u_0$ of the monomer as $U_B(0) = -\epsilon u_0$, where $k_I = \epsilon c^2 k$, and $c = \frac{\theta_S}{2q_c}$. The full binding potential can then be expressed as follows (see Supp. Fig. 3

$$U_B(q) = H(q_c - q)\ \epsilon u_0 \left( \frac{q^2}{q_c^2} - 1 \right). \tag{7}$$

###### 1.2.1 The inter-site distances

The distance between binding sites of a monomer in $S$-form and polymer in $P$-form depends on the geometrical properties of the monomers in a non-trivial manner. Therefore, to study the binding of two monomers we consider that building blocks only have two degrees of freedom:

- The angle $\theta$, which describes the changes in monomer geometry.

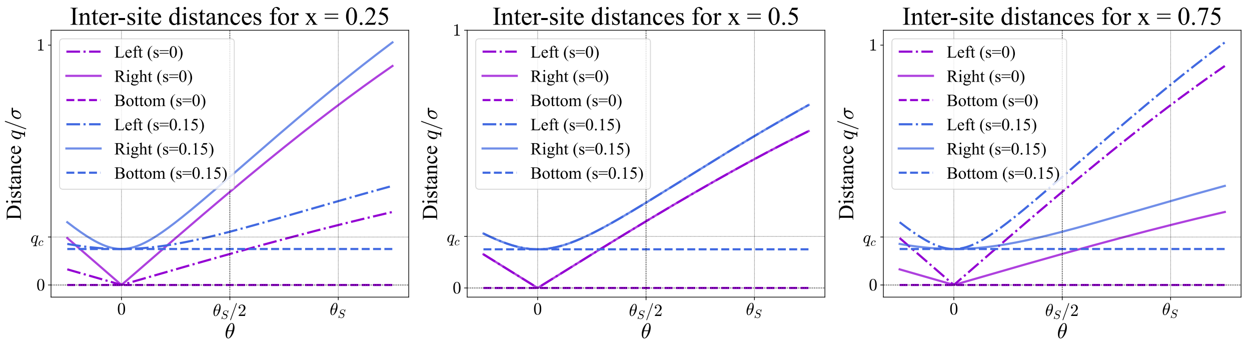

Supplementary Figure 5: Inter-site distance $q$ as a function of $s$ and $\theta$ for three values of $x$. Here $q_c = 0.25\sigma$ and $\theta_S = \pi/3$. Note that for $x = 0.25$ and $x = 0.75$ the two top binding sites cross the cutoff distance $q_c$ on opposite sides of the internal energy landscape barrier (at $\theta_S/2$).

- The inter-base separation $s$, which accounts for the one-dimensional motion of the monomers along the *base* axis (see Fig. 3B in the Main Text and Supp. Fig. 4).

In this scenario, the inter-site distances forming the left (L), right (R) interfaces, and the distance between bottom-binding pairs is given by

$$q_L^2(\theta, s) = s^2 + 2(1-\cos\theta)x(x+s) \tag{8}$$

$$q_R^2(\theta, s) = s^2 + 2(1-\cos\theta)(1-x)(1-x+s) \tag{9}$$

$$q_B^2(\theta, s) = s^2 \tag{10}$$

In general, for any opening angle $\theta$ and separation $s$, $q_L < q_R \iff x < 0.5$, $q_L = q_R \iff x = 0.5$ and $q_L > q_R \iff x > 0.5$ as shown in Suppl. Fig. 5.

##### 1.2.2 The binding energy landscape

We now study how the energy landscape of the system changes when two monomers bind, by considering the following scenarios:

- Case 1: A $S$-form monomer binding a $P$-form seed monomer from the left.
- Case 2: A $S$-form monomer binding a $P$-form seed monomer from the right.
- Case 3: A $S$-form monomer binding two $P$-form monomers (polymer core)

To define the notation, we take the $S$-form monomer as the reference and in what follows, denote case 1 (2) by $R$ ($L$), as it accounts for the formation of an interface on the right (left) side of the monomer; Case 3 is denoted by $M$. The energy landscape associated to each case, as a function of the opening angle $\theta$ and the building block separation $s$, can then be written as follows:

$$U_L(\theta,s) = u_0 \left[ H\left(\frac{\theta_S}{2}-\theta\right)\kappa\left(\frac{4\theta^2}{\theta_S^2}-1\right) + H\left(\theta-\frac{\theta_S}{2}\right)\left(\frac{4(\theta-\theta_S)^2}{\theta_S^2}-1\right) + H\left(q_c - q_L(\theta,s)\right)\epsilon\left(\frac{q_L(\theta,s)^2}{q_c^2}-1\right)\right] \tag{11}$$

$$U_R(\theta,s) = u_0 \left[ H\left(\frac{\theta_S}{2}-\theta\right)\kappa\left(\frac{4\theta^2}{\theta_S^2}-1\right) + H\left(\theta-\frac{\theta_S}{2}\right)\left(\frac{4(\theta-\theta_S)^2}{\theta_S^2}-1\right) + H\left(q_c - q_R(\theta,s)\right)\epsilon\left(\frac{q_R(\theta,s)^2}{q_c^2}-1\right)\right] \tag{12}$$

$$U_M(\theta,s) = u_0 \left[ H\left(\frac{\theta_S}{2}-\theta\right)\kappa\left(\frac{4\theta^2}{\theta_S^2}-1\right) + H\left(\theta-\frac{\theta_S}{2}\right)\left(\frac{4(\theta-\theta_S)^2}{\theta_S^2}-1\right) + H\left(q_c - q_L(\theta,s)\right)\epsilon\left(\frac{q_L(\theta,s)^2}{q_c^2}-1\right) + H\left(q_c - q_R(\theta,s)\right)\epsilon\left(\frac{q_R(\theta,s)^2}{q_c^2}-1\right)\right] \tag{13}$$

In general, as the equations above show, the assembly/disassembly kinetics will be different on either side when $q_L(\theta,s) \neq q_R(\theta,s)$, since the dynamics of the building block will evolve along different energy landscapes; to see this, it suffices to compare $U_L(\theta,s)$ and $U_R(\theta,s)$ above. Therefore, we can expect asymmetric assembly kinetics when the building block contains a hinge asymmetrically coupling the two interfaces, i.e., $x \neq 1/2$.

##### 1.3 Conditions for polymer stability

To study stability of the polymer, we consider the case where $s = 0$, i.e., where the monomer is bound to the polymer by its bottom binding site (see section 1.2.1). For the interface to fully form, the monomer must switch conformations and close such that the upper binding sites can also interact. Defining $f(\theta) = \sqrt{2\,(1-\cos(\theta))}$, the polymerisation potentials in Eqs.11-13 can be expressed as

$$U_L(\theta,s=0) = u_0 \left[ H\left(\frac{\theta_S}{2}-\theta\right)\kappa\left(\frac{4\theta^2}{\theta_S^2}-1\right) + H\left(\theta-\frac{\theta_S}{2}\right)\left(\frac{4(\theta-\theta_S)^2}{\theta_S^2}-1\right) + H\left(q_c - f(\theta)x\right)\epsilon\left(\frac{f(\theta)^2x^2}{q_c^2}-1\right) - \epsilon\right] \tag{14}$$

$$U_R(\theta,s=0) = u_0 \left[ H\left(\frac{\theta_S}{2}-\theta\right)\kappa\left(\frac{4\theta^2}{\theta_S^2}-1\right) + H\left(\theta-\frac{\theta_S}{2}\right)\left(\frac{4(\theta-\theta_S)^2}{\theta_S^2}-1\right) + H\left(q_c - f(\theta)(1-x)\right)\epsilon\left(\frac{f(\theta)^2(1-x)^2}{q_c^2}-1\right) - \epsilon\right] \tag{15}$$

$$U_M(\theta,s=0) = u_0 \left[ H\left(\frac{\theta_S}{2}-\theta\right)\kappa\left(\frac{4\theta^2}{\theta_S^2}-1\right) + H\left(\theta-\frac{\theta_S}{2}\right)\left(\frac{4(\theta-\theta_S)^2}{\theta_S^2}-1\right) + H\left(q_c - f(\theta)x\right)\epsilon\left(\frac{f(\theta)^2x^2}{q_c^2}-1\right) + H\left(q_c - f(\theta)(1-x)\right)\epsilon\left(\frac{f(\theta)^2(1-x)^2}{q_c^2}-1\right) - 2\epsilon\right] \tag{16}$$

We will now explore the self-assembly kinetics resulting from these energy landscapes. In particular, we are interested in the existence and values of well-defined minima for $\theta = 0$ and $\theta = \theta_S$ and in the activation barriers for switching between the two, which will control the kinetic rates of binding and unbinding (i.e., forming or breaking the full interface).
The minima of the potentials above for the polymer form, namely, for $\theta = \theta_P = 0$ is

$$U_L^P = U_L(0,0) = -u_0(\kappa + 2\epsilon) \tag{17}$$
$$U_R^P = U_R(0,0) = -u_0(\kappa + 2\epsilon) \tag{18}$$
$$U_M^P = U_M(0,0) = -u_0(\kappa + 4\epsilon) \tag{19}$$

indicating that $\epsilon$ can stabilise the bound $P$-form of the building block.
The solution configuration minima depend on $q_c$, $f_S = f(\theta_S)$ and $x$,

$$U_L^S = U_L(\theta_S,0) = -u_0\left[1+\epsilon\left(1-H(q_c - f_S x)\left(\frac{f_S^2 x^2}{q_c^2}-1\right)\right)\right]$$
$$U_R^S = U_R(\theta_S,0) = -u_0\left[1+\epsilon\left(1-H(q_c - f_S(1-x))\left(\frac{f_S^2(1-x)^2}{q_c^2}-1\right)\right)\right]$$
$$U_M^S = U_M(\theta_S,0) = -u_0\left[1+\epsilon\left(2-H(q_c - f_S x)\left(\frac{f_S^2 x^2}{q_c^2}-1\right)-H(q_c - f_S(1-x))\left(\frac{f_S^2(1-x)^2}{q_c^2}-1\right)\right)\right]$$

At the molecular scale, interaction ranges are typically quite short. We can therefore assume $q_c \ll \sigma$. If $q_c < f_S \min(x, 1-x)$ such that in the solution configuration the top binding sites escape the binding potential on both sides even at $s = 0$, the $S$-form minima become

$$U_L^S = U_L(\theta_S,0) = -u_0(1+\epsilon) \tag{20}$$
$$U_R^S = U_R(\theta_S,0) = -u_0(1+\epsilon) \tag{21}$$
$$U_M^S = U_M(\theta_S,0) = -u_0(1+2\epsilon) \tag{22}$$

Finally, we evaluate the stability of the $P$-form monomer when bound to a polymer in each scenario by computing the energy difference between the two minima:

$$\Delta U_L = U_L^P - U_L^S = u_0(1-\kappa-\epsilon) \tag{23}$$
$$\Delta U_R = U_R^P - U_R^S = u_0(1-\kappa-\epsilon) \tag{24}$$
$$\Delta U_M = U_M^P - U_M^S = u_0(1-\kappa-2\epsilon) \tag{25}$$

Note that the energy difference is the same for left (L) and right (R) interface formation, as the two minima coincide. Crucially, the energy difference for the polymer core differs with that of the left and right interfaces: it is harder for monomer to switch from its $P$-form to its $S$-form within the core than on the edges, because it is stabilised by two interfaces instead of a single one.
To summarize, from the analysis above we can extract that polymer edges are stable if and only if $\epsilon > 1-\kappa$, and that the polymer core is stable if and only if $\epsilon > \frac{1-\kappa}{2}$. These conditions can be

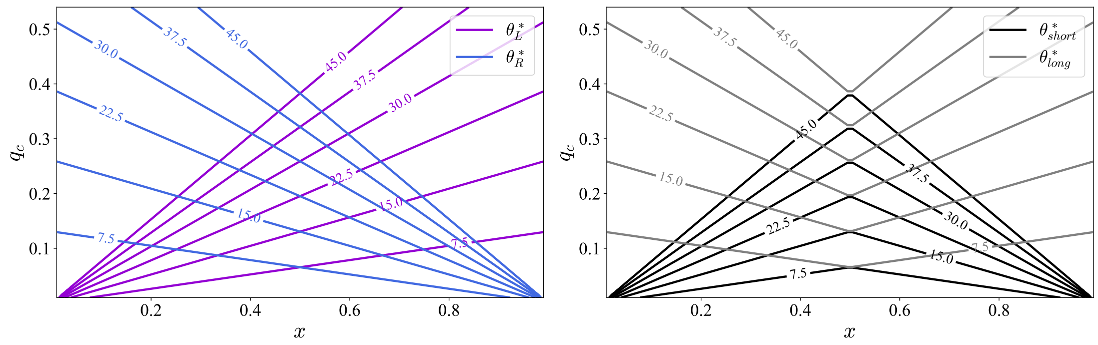

Supplementary Figure 6: a) $\theta^*_L$ and $\theta^*_R$ as a function of $x$ and $q_c$. b) $\theta^*$ on the short and long side of the building block as a function of $x$ and $q_c$. All angles are in degrees.

translated into expected growth or disassembly regimes as follows:

$$0 < \epsilon < \frac{1-\kappa}{2} \iff \text{the polymer tends to fragment} \quad (26)$$

$$\frac{1-\kappa}{2} < \epsilon < 1-\kappa \iff \text{the polymer tends to shrink from its edges} \quad (27)$$

$$1-\kappa < \epsilon \iff \text{the polymer tends to grow} \quad (28)$$

Such conditions delimit the regions shown in Fig. 4B in the main text.

#### 1.4 Catalyzed interface formation

The self-assembly kinetics of the monomers will be controlled by the combination of their internal potential $U_I(\theta)$ (see Eq. 1), which controls fluctuations in $\theta$, and by the the binding potential $U_{L,R}(q)$ describing the formation of the specific interface (see Eqs. 11 and 12). We define as $\theta^*_L$ and $\theta^*_R$ the opening angles for which the binding site distances $q_L = q_c$ and $q_R = q_c$ respectively. These denote the points on the fluctuation energy landscape for which the binding contribution *deforms* the landscape, and satisfy:

$$2(1-\cos\theta^*_L)x^2 + s^2 = q_c^2 \qquad \text{and} \qquad 2(1-\cos\theta^*_R)(1-x)^2 + s^2 = q_c^2 \quad (29)$$

which we can solve to obtain

$$\theta^*_L = \arccos\left(1 - \frac{q_c^2 - s^2}{2x^2}\right) \qquad \text{and} \qquad \theta^*_R = \arccos\left(1 - \frac{q_c^2 - s^2}{2(1-x)^2}\right). \quad (30)$$

For $s = 0$, the expressions reduce to

$$\theta^*_L = \arccos\left(1 - \frac{q_c^2}{2x^2}\right) \qquad \text{and} \qquad \theta^*_R = \arccos\left(1 - \frac{q_c^2}{2(1-x)^2}\right). \quad (31)$$

In general, $\theta^*_L \neq \theta^*_R$, except when $x = 1/2$ (see Suppl. Fig. 6). This feature is at the root of the asymmetric binding kinetics of our toy model, and allows us to distinguish three different scenarios depending on $x < 1/2$ and $q_c$ (see Suppl. Fig. 7):

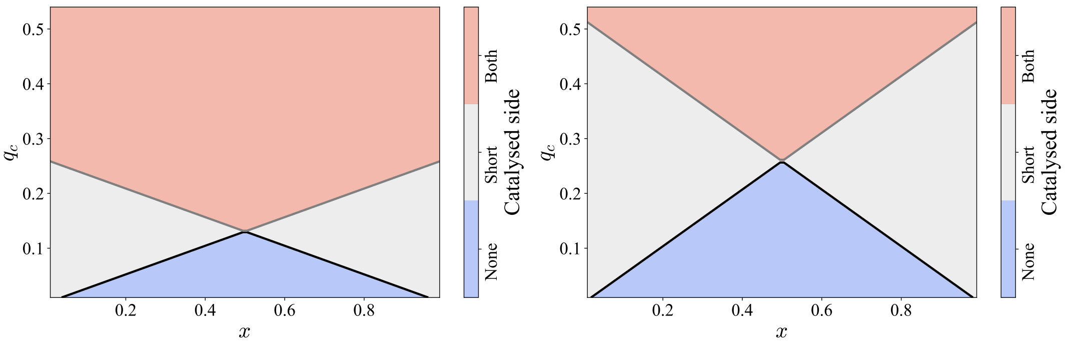

Supplementary Figure 7: Closing reaction catalysis map. Diagram indicating on which side the closing reaction is catalysed by the presence of a polymer at distance $s = 0$ for $\theta_S = 30$ degrees (A) and $\theta_S = 60$ degrees (B).

- $S$-to-$P$ switch is not catalysed on either side if $\theta^* < \theta_S/2$ on both sides.
- $S$-to-$P$ switch is catalysed on the short side but not the long side if $\theta_R^* < \theta_S/2 < \theta_L^*$.
- $S$-to-$P$ switch is catalysed on both sides if $\theta_R^* \leq \theta_L^* < \theta_S/2$.

To understand the kinetics of self-assembly, we study the barriers separating two minima: the $P$-form when the building block binds to a polymer seed, and the $S$-form of a freely diffusing monomer. We consider two different scenarios (Fig. 8):

- **No catalysis**: If $\theta^* < \theta_S/2$ then the internal energy landscape remains unchanged in the interval $[\theta_S/2, \theta_S]$ (except for global shifts affecting all $\theta$ values which result from pure $s$ contributions). This implies that the closing barrier will remain the same as for the free monomer ($\Delta U^\ddagger = u_0$) and the opening barrier will be controlled by the new $P$ form minimum: $\Delta U^\dagger = (\kappa + \epsilon)u_0$ for monomers binding only on one side (left or right), or $\Delta U^\dagger = (\kappa + 2\epsilon)u_0$ for monomers binding on both sides (left and right, so two binding sites are contributing $\epsilon u_0$ when closed).
- **Catalysis**: If $\theta^* > \theta_S/2$ then the internal energy landscape will be deformed at its barrier, decreasing it. This can result in a number of different scenarios depending on the specific values of the parameters and particularly on how strong the binding attraction is. Let us explore this in more detail below.

##### 1.4.1 Catalysis: how the switching barrier is changed by the binding interaction

When $\theta^* > \theta_S/2$ then, because all our potentials are harmonic, the full potential in the interval $\theta \in [\theta_S/2, \theta^*]$ will be an upwards parabola centred at some minimum $\theta_m \in [\theta_S/2, \theta^*]$ which will depend on the parameters of the potentials combined. Note here that, more generally, for an internal landscape that monotonically decreases in that interval and a binding potential that monotonically increases in that interval, the resulting full potential (from the sum of the two) will also have a single minimum in the closed interval $[\theta_S/2, \theta^*]$. Three scenarios are possible then:

1. The minimum of the full potential in the interval $[\theta_S/2, \theta^*]$ corresponds to $\theta_m = \theta_S/2$, such that the potential is monotonically increasing in the interval.

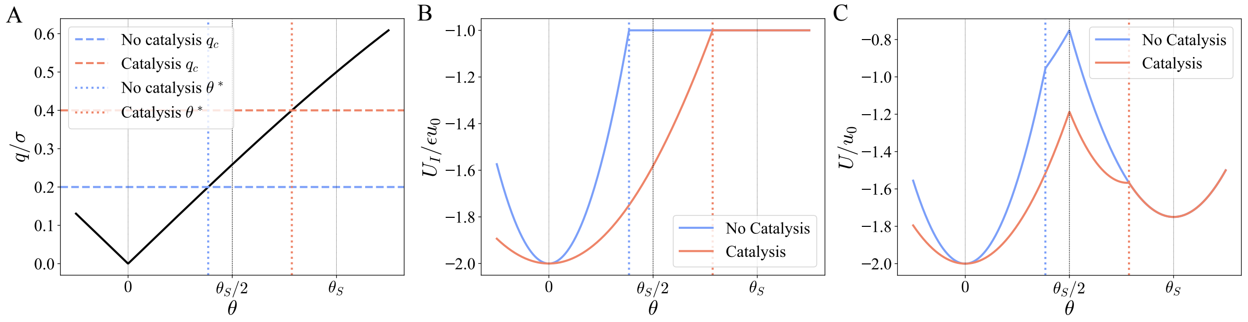

Supplementary Figure 8: A) Distance between the top binding sites of a symmetric building block at a distance $s = 0$ from a polymer as a function of the opening angle $\theta$ for $q_c = 0.4\sigma$ (catalysis, red) and $q_c = 0.2\sigma$ (no catalysis, blue). Vertical dotted lines indicate the critical opening angle $\theta^*$ for which $q(\theta) = q_c$ in each case. B) Binding sites interaction potential as a function of $\theta$ for the case displayed in A. The binding potential is non-zero at the barrier $\theta_S/2$ only for the catalysis scenario. C) Full energy landscape of the switch (internal and binding potentials) for the two scenarios. The barrier for switching is lower for the red curve (catalysis). Parameters: $x = 0.5$, $\theta_S = 60$ degrees, $\kappa = 0.5$, $\epsilon = 0.75$.

This will only happen for large values of $\epsilon$, such that the binding potential overcomes the internal potential barrier. While a precise value of the critical binding strength required for this regime is not tractable, we can approximate it (using a Taylor expansion of $q$) to $\epsilon\mu^2 > 1$, where $\mu = \frac{\theta_S m}{2q_c}$.

In this regime the activation barrier for binding/unbinding thus lies at $\theta^*$ and the barrier will be controlled by the value of the potential at that point.

Conveniently, this point lies in the purely internal part of the potential and therefore does not depend on $\epsilon$. Given that in the $S$ form minimum $U_I(\theta_S) = -u_0$, we can define $\lambda = U_I(\theta^*) - U_I(\theta_S)$ the difference between the two points of interest, such that $U_I(\theta^*) = u_0(\lambda - 1)$. For our choice of potentials $\lambda = \frac{4(\theta^* - \theta_S)^2}{theta_S^2}$ but this description holds for any choice of potential.

In this scenario then, the closing barrier will be given by $\Delta U^\ddagger = \lambda u_0$ and the opening one will either be $\Delta U^\dagger = (\kappa + \epsilon + \lambda - 1)u_0$ for monomers binding only on one side (left or right), or $\Delta U^\dagger = (\kappa + 2\epsilon + \lambda - 1)u_0$ for monomers binding on both sides (left and right, so two binding sites are contributing $\epsilon u_0$ when closed).

2. The minimum of the full potential in the interval corresponds to $\theta_m = \theta^*$, such that the potential is monotonically decreasing in the interval.

   This will only happen for small values of $\epsilon$, such that the internal barrier overcomes the binding strength. Again a precise computation is difficult but in our model we can deduce the following condition on the binding strength: $\epsilon\mu^2 - 2\mu + 1 < 0$, which only holds if $\epsilon < 1$.

   In this regime the activation barrier for binding/unbinding thus lies at $\theta_S/2$ and the barrier will be controlled by the contributions of the binding potential to that point.

   We can express the contribution of the moving binding site to the full potential at the barrier $\theta_S/2$ as $U^\dagger = -\gamma\epsilon u_0$, with $\gamma \in [0, 1]$ representing the fraction of the attraction *felt* by the

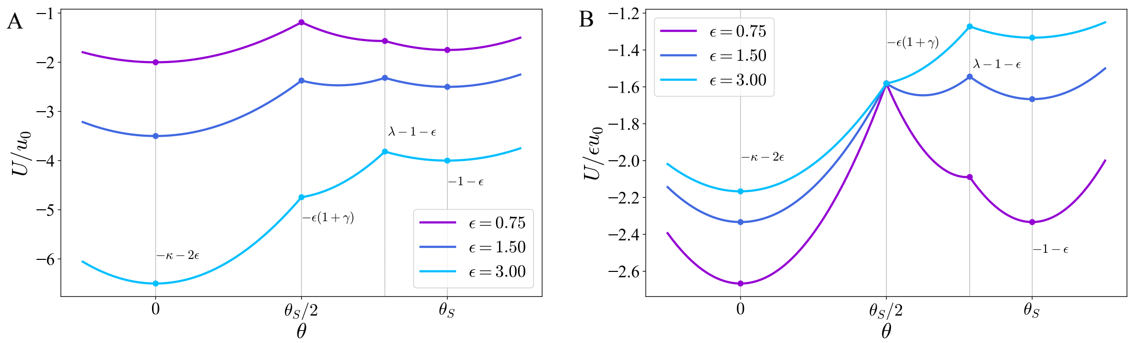

Supplementary Figure 9: A) Full energy landscape of the switch (internal and binding potentials) for a symmetric building block ($x = 0.5$) at a distance $s = 0$ from a polymer (bound by the bottom) in the catalysis regime ($\theta_S = 60$ degrees, $q_c = 0.4\sigma$). The internal potential is given by $\kappa = 0.5$ and three binding regimes are shown: soft ($\epsilon = 0.75$), intermediate ($\epsilon = 1.50$) and strong ($\epsilon = 3.00$). In all cases the switching barrier is affected, but the resulting energy landscape is qualitatively different for stronger binding. B) Same energy landscapes as in A, but normalised by $\epsilon$ to illustrate the qualitative difference in the landscape shape.

moving binding site at the barrier point. For our choice of binding potential and model we find that $\gamma_L = \max(1 - \frac{x^2}{q_c^2} 2(1 - \cos\theta_S/2), 0)$ and $\gamma_R = \max(1 - \frac{(1-x)^2}{q_c^2} 2(1 - \cos\theta_S/2), 0)$, which becomes $\gamma = 0$ when $\theta^* < \theta_S/2$. Importantly, the use of $\gamma$ is general to any choice of binding potential that is monotonically increasing in the interval $q \in [0, q_c]$ from $-\epsilon u_0$ to 0.

In this scenario then, for a monomer binding only from one side, the closing barrier will be given by $\Delta U^{\ddagger} = (1 - \epsilon\gamma) u_0$ and the opening barrier by $\Delta U^{\dagger} = (\kappa + \epsilon(1 - \gamma)) u_0$. If the monomer is binding on both sides instead, then the barriers become $\Delta U^{\ddagger} = (1 - \epsilon(\gamma_L + \gamma_R)) u_0$ and $\Delta U^{\dagger} = (\kappa + \epsilon(1 - \gamma_L - \gamma_R)) u_0$.

3. The minimum lies somewhere in the interval $\theta_m \in (\theta_S/2, \theta^*)$, such that a third metastable state emerges in the opening/closing pathway due to the presence of a catalysing polymer at $s = 0$. Understanding the kinetics of this regime becomes more complicated due to the emergence of this third minimum at $\theta = \theta_m$, but this only happens for moderate values of $\epsilon$, where the contribution of the binding is comparable to the internal switching barrier. In general we will neglect this regime for simplicity.

Overall then, the switching energy landscape resulting from the catalysis of the reaction will depend on the strength of the binding potential, decreasing the barrier proportionally to $\epsilon$ until saturation when the barrier switches from $\theta_S/2$ to $\theta^*$ for strong binding. Note that, this will be the dominant regime of interest for polymerisation, where we need a strong enough $\epsilon$ to stabilise the closed form of the monomer in the polymer.

In Figure 9 we illustrate these three scenarios for a symmetric model building block ($x = 0.5$) with $\theta_S = 60$ degrees and $\kappa = 0.5$ subject to catalysis ($q_c = 0.4\sigma$ such that $\theta^* > \theta_S/2$). For soft binding (purple curve, $\epsilon = 0.75$) the barrier is still at $\theta_S/2$ and only minimally decreased. For intermediate binding (dark blue curve, $\epsilon = 1.50$) a third minimum emerges for $\theta \in (\theta_S/2, \theta^*)$. For strong binding (light blue curve, $\epsilon = 3.00$), the barrier at $\theta_S/2$ disappears and the kinetics are controlled by a new activation barrier at $\theta^*$.

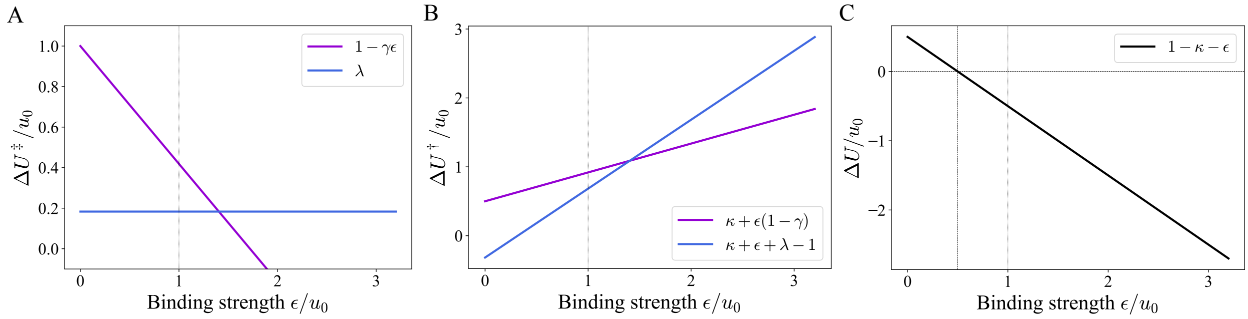

Supplementary Figure 10: A) Switching barrier for the solution-to-polymer transition as a function of $\epsilon$. B) Switching barrier for the polymer-to-solution transition as a function of $\epsilon$. C) Energy difference between the two configurations ($\Delta U = U_P - U_S$) as a function of $\epsilon$. The $P$ form becomes more stable for $\epsilon > 1-\kappa$. In all panels we work with a symmetric model building block ($x = 0.5$) with $\theta_S = 60$ degrees, $q_c = 0.4\sigma$ and $\kappa = 0.5$, resulting in $\lambda \sim 0.18$ and $\gamma \sim 0.58$.

From this analysis we can approximate the value of the switching barrier (solution-to-polymer transition) for a polymer binding on a single side as a function of $\epsilon$ (Fig. 10A):
- In the soft binding limit, $\epsilon << 1$, the barrier is controlled by $\epsilon$ and given by the following expression:

$$\Delta U^{\ddagger} = (1-\epsilon\gamma)u_0$$

- In the strong binding limit, $\epsilon >> 1$, the barrier is independent of $\epsilon$ and given by the following expression:

$$\Delta U^{\ddagger} = \lambda u_0$$

Similarly, we can also estimate the value of the polymer-to-solution transition barrier for a polymer binding on a single side as a function of $\epsilon$ (Fig. 10B):
- In the soft binding limit, $\epsilon << 1$, the barrier is controlled by $\epsilon$ and given by the following expression:

$$\Delta U^{\dagger} = (\kappa + \epsilon(1-\gamma))\, u_0$$

- In the strong binding limit, $\epsilon >> 1$, the barrier is independent of $\epsilon$ and given by the following expression:

$$\Delta U^{\dagger} = (\kappa + \epsilon + \lambda - 1)\, u_0$$

Importantly, the energy difference between the $S$ and $P$ configurations remains the same regardless of the regime (Fig. 10C):

$$\Delta U = 1 - \kappa - \epsilon$$

##### 1.5 Binding and unbinding kinetics for $s = 0$

Let us now go back to our original problem and investigate how these catalysis scenarios can translate into polar self-assembly kinetics. So far in our discussion of how a polymer at distance $s = 0$ can catalyse the switch reaction we have neglected the role of $x$, sticking to symmetric building blocks throughout. However, we have seen before that $x$ plays a crucial role in how the binding interactions affect the internal switching energy landscape. We will now explore the difference in

switching barriers between binding on the left or on the right as a function of the building block geometry.
For the sake of simplicity we will limit ourselves to the case of strong binding ($\epsilon >> 1$), which is most relevant to polymerisation and stable protofilament formation. As shown just above, in this scenario the binding interaction switches the barrier from $\theta_S/2$ to $\theta^*$ (where the binding site starts to *feel* the interaction) if $\theta^* > \theta_S/2$ (catalysis regime).
In this scenario, depending on the values of $x$, $q_c$ and $\theta_S$, we distinguish different kinetic regimes:

- **No catalysis on either side**: $\theta^* < \theta_S/2$ on both sides and the closing reaction is not catalysed on either side of the building block. The switching barrier is unaffected by the presence of the polymer at distance $s = 0$ and we expect **non polar polymerisation**. This corresponds to the dark blue triangle below the black dashed line in Fig. 11.
- **Only the short side is catalysed**: $\theta^*_{long} < \theta_S/2 < \theta^*_{short}$ such that the switching barrier is lowered on the short side but unaffected on the long side, resulting in different kinetics of self-assembly. We expect **polar polymerisation** with **faster kinetics on the short side**. This is illustrated in Fig. 11 by the increase in $\Delta\Delta U^{\ddagger}$ as the building block becomes more asymmetric, between the black, red and gray dashed lines.
- **Both sides are catalysed**: $\theta_S/2 < \theta^*_{long} < \theta^*_{short}$ such that the switching barrier is lowered on both sides, but it is lowered more on the short side, resulting again in a difference in switching barriers. We expect **polar polymerisation** with **faster kinetics on the short side**. This is illustrated in Fig. 11 by the increase in $\Delta\Delta U^{\ddagger}$ as the building block becomes more asymmetric above the gray and below the red dashed lines.
- **Short side loses barrier**: for $\theta^*_{short} > \theta_S$ (above the red dashed line in Fig. 11) the short side cannot escape the binding potential for $s = 0$. Here the $S$ form minimum disappears and the kinetics are ill defined. We expect a quick closing of the building block. In the light red region in Fig. 11 the long side is not catalysed, such that we expect **polar polymerisation** with **faster kinetics on the short side**, and in the purple region the long side becomes catalysed with decreasing barrier as we approach the blue dashed line. Here the kinetics are ill defined.
- **Both sides lose barriers**: for $\theta_S < \theta^*_{long} < \theta^*_{short}$ (light blue region in Fig. 11) neither side can escape the binding potential for $s = 0$. The $S$ form minimum disappears on both sides and the kinetics become ill defined again, but we expect **non polar polymerisation** as there are no activation barriers for closing on either side.

### 2 LAMMPS implementation of the elastic network model

In this subsection, we detail the implementation of the minimal elastic network model in the Molecular Dynamics (MD) simulation software LAMMPS, as well as the simulation set-up used to evaluate the emergence of asymmetric kinetics and treadmilling dynamics. Codes to reproduce the results are provided as a repository.

#### 2.1 Particle-based implementation of elastic network

The particle-based representation of the elastic network model in section 1.1 consists of 7 different types of spherical particles with diameter $\sigma$, which sets the length scale of the simulation. The

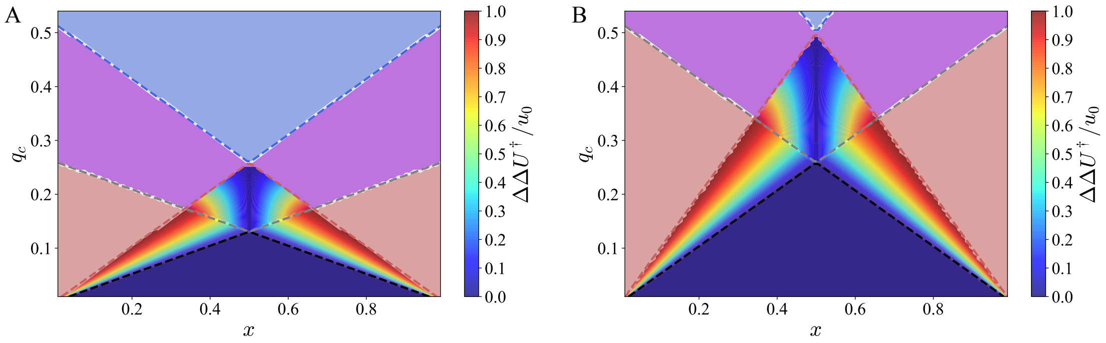

Supplementary Figure 11: Closing barriers difference map as a function of $x$ and $q_c$ for building blocks binding to a polymer at distance $s=0$ on the left and right side. $\Delta\Delta U^\ddagger = \Delta U^\ddagger_{long} - \Delta U^\ddagger_{short}$, hence why the barrier difference is symmetric around $x=1/2$. We consider only the strong binding regime ($\epsilon \gg 1$) and illustrate the emerging asymmetry in the energy barriers between the two sides for $\theta_S = 30$ degrees (A) and $\theta_S = 60$ degrees (B). In the red, purple and blue shaded regions the barrier difference cannot be computed (see main text), but for the red and purple regions we expect faster kinetics on the short side while we expect symmetric kinetics in the blue region.

| **Bond properties** | | |
|---|---|---|
| Bond type | Stiffness | Equilibrium length |
| 1 | $10^5$ | $x \times L_S$ |
| 2 | $10^5$ | $(1-x) \times L_S$ |
| 3 | $10^5$ | $L_S$ |
| **Angle properties** | | |
| Angle type | Stiffness | Equilibrium angle (rad) |
| 1 | $10^5$ | $\pi$ |
| 2 | $10^5$ | $\pi/2$ |
| 3 | $10^5$ | $\theta_P = \pi/2$ |
| 4 | $10^5$ | $\theta_S - \theta_P$ |

Table 1: List of the parameters used to simulate the elastic network model in LAMMPS.

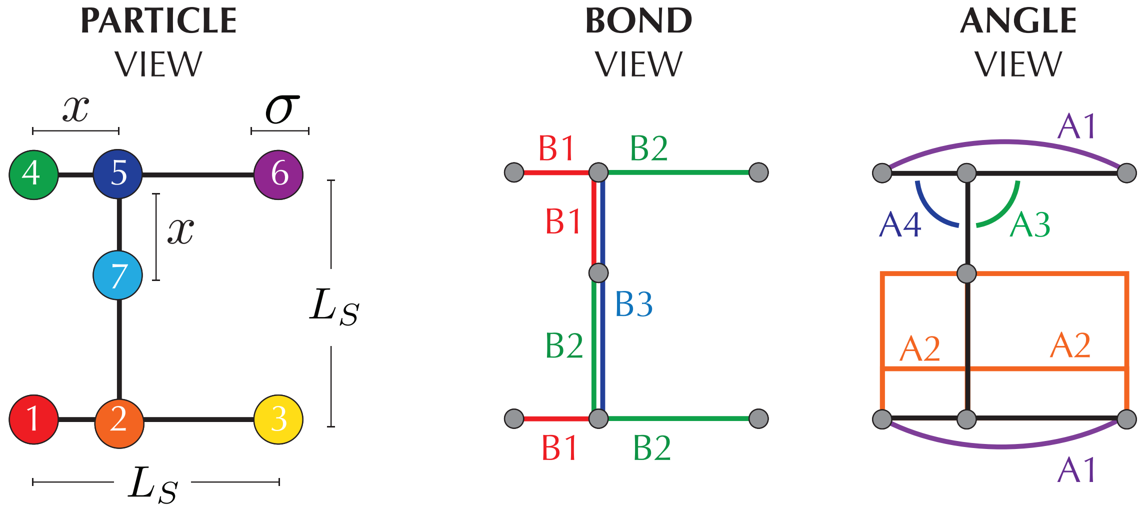

Supplementary Figure 12: Particle-based realization of the elastic network model in LAMMPS. To reproduce the geometrical features of the elastic network, we use a building block made of 7 types of particles (particle view), 3 different types of bonds (bond view) and 4 types of angles. Particle, bond and angle properties are detailed in subsection 2.1.

overall scale of the building block is given by $L_S = 10\sigma$ (see Supplementary Fig. 12). Particles 1, 3, 4 and 6 correspond to binding sites in the building block, while 2, 5 and 7 play structural roles to maintain the monomer scaffold in simulations. In particular, particle 7 prevents the inversion of the hinging arm (see Fig. 2 in the main text). Particles 1-2, 4-5, and 5-7 are connected by harmonic bonds of type 1, while particles 2-3, 5-6 and 2-7 are connected by harmonic bonds of type 2; lastly, particles 2-5 are connected by a harmonic bond of type 3. Additionally, we consider angle potentials of type 1 between particles 1-2-3 and 4-5-6, of type 2 between particles 1-2-5, 5-2-3, 1-2-7 and 7-2-3, of type 3 between 2-5-6 and of type 4 between 4-5-2. Angles of type 3 and 4 control the internal energy landscape of the monomer. To allow for their breaking and formation at cutoff $\theta_c = \theta_S/2$, we have developed a new angle style for LAMMPS (see *harmswitch* potential in the repository). Parameters for each bond and angle types can be found in Table 1.
We conduct two-dimensional simulations using the LAMMPS enforce2d fix. Particles of type 1, 2 and 3 in every monomer are constrained by a trapping force to move along the $y = 0$ line. We use $dt = 10^{-2}$ as the simulation time step. Interactions between binding sites of type 4 and 6 in different monomers are mediated by a cosine squared potential with strength $\varepsilon$ and cutoff $r_c$; interactions between binding sites of type 1 and 3 are mediated by a WCA potential with cutoff $\sigma$. Within the same monomer, particles of type 4 and 7 repel each other via a WCA potential when $r < \sigma$.

#### 2.2 Simulations for asymmetric polymerization

To test for asymmetric polymerisation, we fix a monomer in the middle of a rectangular simulation box with reflecting walls. This monomer, which will act as a polymerisation seed and is initialized in the $P$-form, is not time-integrated during the simulation. We then add two monomers at either side of the seed and time-integrate their dynamics to study the growth of left and right ends of the polymer. To keep the concentration in the left- and rightmost ends of the polymer constant as the filament grows, we alternate between 50-step-long MD runs and evaluations of the state of the system using a python script. The script analyses the number of connected components in the system by considering each monomer as a node, and each interaction between monomers as an edge. Two monomers are considered to be interacting if their top binding sites (particles of type 4 and 6) are within a distance $r < 0.9r_c$. When the number of monomers at a given end of the polymer decreases due to a binding event, we increase the box size by moving the location of the simulation wall at the corresponding side, and add a new monomer in the $S$-form which is then allowed to diffuse. The right (left) wall is displaced a factor $\delta = 2 \times L_S$ in the +(-)x direction after each binding event. Freshly added monomers are initialised at a fixed distance from the corresponding polymer interface equal to $1.3 \times L_S$. To characterize the kinetic asymmetry of the system, we track the center of mass (COM) dynamics of the growing polymer, as well as the positions of the left- and right-most monomers in the filament, which, as the seed of the filament is fixed in space, allow us to quantify the kinetic polarity of the system.

##### 2.3 Treadmilling simulations

Simulations to study treadmilling rely on the same set-up as the one detailed to study asymmetric polymerisation, i.e., the number of monomers at either side of the polymer is also kept constant. However, in this case all monomers are free to move. The system is initialized as a filament with $N = 25$ monomers bound in the $P$-form, and monomers are added to either end of the filament to allow for polymerisation. To introduce energy dissipation in the system, we randomly choose monomers within the polymer for *hydrolysis* every 50 steps in the simulation. Once a monomer is hydrolysed, the binding strength of both of its top binding sites (particles of type 4 and 6) is decreased from $\varepsilon$ to $0 < \varepsilon_H < \varepsilon$. The final strength $\varepsilon_H$ is governed by the fragmentation probability of the filament, and ensures (i) that the filament will not break apart due to hydrolysis and (ii) that detachment will predominantly happen on one end of the filament due to thermal fluctuations.
When a monomer detaches, we remove it from the system, and replenish the monomer pool with fresh, unhydrolysed monomers following the procedure detailed in subsection 2.2. As we are interested in single filament dynamics, if the filament fragments, we keep the longest fragment and delete the rest of the monomers from the system. The simulation then continues with the same constant concentration procedure.

#### 3 LAMMPS implementation of the colloidal design

In this section, we will provide further details on the MD simulations ran using the colloidal monomeric design. Sample code for all of the simulations used is in the provided repository.

##### 3.1 Colloidal monomeric design

The full particle and bond implementation is shown in Supplementary Fig 13. The particles are of diameter $\sigma$ and the bond equilibrium length is $L = \sigma$, while the stiffness is $10^3 k_B T$ (here the equilibrium bond length is shown as larger to visualise the bonds and particles together). The missing bond between particles 2-7 is allowing for the conformational switch between 2 states, which are the only two stable states of the monomer structure.

The double well potential is enforced by interactions between types 3-4 ($\varepsilon_P$) and types 2-7 ($\varepsilon_S$). Interactions between monomers to form a filament are set between types 1-2 and 6-7 ($\varepsilon$). The rest of the pairwise interactions between different types of particles interact simply via volume exclusion (cutoff $= \sigma$). All of the interactions are short range interactions (cutoff $=1.2\sigma$) implemented in LAMMPS using the cosine/squared potential. As before, all of the simulations are run in 2d, using the enforce2d fix in LAMMPS, with a timestep of $dt = 10^{-2}$.

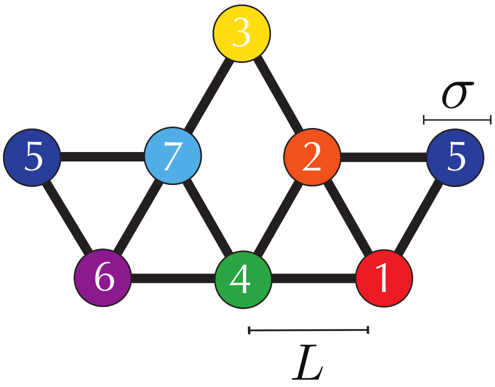

Supplementary Figure 13: Particle-based realization of the colloidal design in LAMMPS. To ensure correct assembly, we use a building block made of 7 types of particles and 1 type of bond.

To study binding kinetics in the colloidal model we develop three sets of simulations:

1. Quantifying binding kinetics from tracking binding events between two monomers
2. Simulations of asymmetric polymerisation and fragmentation
3. Treadmilling simulations

#### 3.2 Tracking binding between two monomers

To probe the effect of the conformational switch we use a simulation setup in which we put two monomers in a square box and track their binding kinetics. One of the monomers has an additional bond between particles 3-4 (see Supplementary Fig 13) and is therefore fixed in the polymeric, self-fitting state. The other monomer corresponds to the design with a conformational switch and acts as a free monomer. This way one monomer acts as a filament end, and the other as the monomer in solution, allowing us to study polymerization kinetics in a simplified setup.
We study the effect of the energetic landscape of the switching monomer on binding kinetics. We run a parameter sweep where we test binding strength of $1 - 10k_BT$ for both $\epsilon_P$ and $\epsilon_S$, while $\epsilon$ is fixed at $10k_BT$. For every value of $\epsilon_S$ and $\epsilon_P$ we perform $10^4$ replicas of binding events where we track first encounter time and full binding time. First encounter time is defined as the time at which one of the particle at the binding site starts interacting with the monomer in polymer form, while full binding corresponds to the whole 3-particle binding site interface forming (interface 1:particles 2,3,7; interface 2:particles 1,4,6 see Supplementary Fig 13).
This gives a distribution of first encounter time and binding time. To make sure to not bias the distribution, we initialise the two monomers randomly in the simulation box, varying the initial distance between the two. Additionally, to get the correct tails of both distributions of less likely

binding events, we only focus on binding either at the pointed or back end of the polymer form monomer. We do this by removing the interactions on one of the binding sites of the solution form monomer, guaranteeing sampling of binding only on one polymer end.

We can then form a distribution of first encounter times and binding times, which we fit to an exponential and gamma distribution, respectfully. An example of such distributions is shown in Fig 14. The binning is done using the Freedman-Diaconis rule and the fitting distributions are chosen based on the data shown. Then we can extract the rate out of the fitted distribution and plot heatmaps of the ratio of the distributions binding on one and on the other polymer end. This corresponds to the heatmap shown in Fig 5B for encounter times showing a clear asymmetry for monomers in solution form.

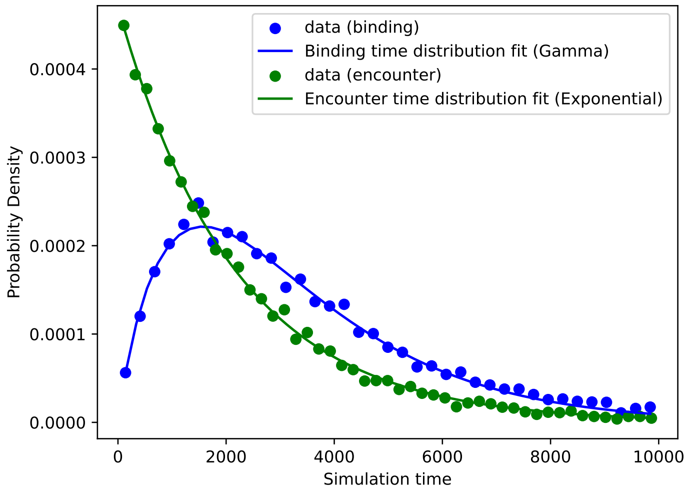

Supplementary Figure 14: Fitting distributions to measured data: Example of binding and encounter times for $\epsilon_P = 3$ kT and $\epsilon_S$=7 kT

##### 3.3 Simulations of asymmetric polymerisation and fragmentation

After establishing how the bistable landscape affects encounter asymmetry, we set the internal landscape at $(\varepsilon_S, \varepsilon_P) = (6,3)$ and $(\varepsilon_S, \varepsilon_P) = (3,6)$. Next, we perform polymerisation analysis in a similar way as introduced above. We start the simulation in a rectangular box of size $20\sigma$x$10\sigma$. Upon the formation of a dimer, we fix the dimer center of mass in the centre of the box and start extending it in the x direction to add new monomers. We keep the monomer concentration on both polymer ends constant by always having one monomer on both sides of the polymer and by keeping a constant distance between the right/leftmost monomer and the box edge ($\Delta L = 20\sigma$). Every 100 steps, we check if there was a binding event on any of the 2 sides of the polymer, and if so we add a new monomer at a constant diatance from the right/leftmost monomer at a distance of $\Delta = 10\sigma$. The binding is very strong, to ensure there are no disassembly events in the process ($\varepsilon = 15k_BT$). We run 300 replicas of these simulations to extract polymerisation rates. We then collect the number of added monomers to both polymer sides over time from all the replicas, and extract the rates and errors by fitting this data to a linear plot. The extracted rates are plotted in

Fig 5B.

For the fragmentation study, we take a filament of a formed length through polymerisation simulations ($N = 50$) and weaken the binding between monomers to see where in the filament the first detachment happens. Similarly, we run 300 replicas of the simulations with lowered $\varepsilon$ until a breaking event happens. We then record whether this happened in the middle or end of the polymer and plot the probability of end disassembly in Fig 5. Similar analysis was conducted for the initial design in Fig 1, but with ($N = 5$) to prove that even in a situation where the number of end and middle interfaces is the same (N=2), there is no preferential end unbinding. Running these simulations initally in a small simulation box surrounding the filament, we noticed that the presence of walls helped stabilise end monomers, so we perform this analysis in a very big box to avoid these effects($2000\sigma$x$2000\sigma$).

##### 3.4 Treadmilling simulations

After establishing a condition for directional polymerisation at $(\varepsilon_S, \varepsilon_P) = (6, 3)$, and finding the optimal binding strength from the end disassembly analysis $\varepsilon_H = 8k_BT$, we incorporate hydrolysis into the previous setup of a growing filament to test for the emergence of treadmilling. The binding strength of non-hydrollysed monomers is set to $\varepsilon = 15k_BT$ Given the comment from the previous section, where a small box was found to stabilise the filament core from breaking, we impose a different condition to preserve constant monomer concentration on both filament ends. We run the simulation in a large box, to avoid biasing the fragmentation, and instead keep one monomer in a circle or radius $R = 8\sigma$ around the end monomers. This preserves the area for monomers to diffuse similar to the one imposed in the previous analysis, allowing us to set the hydrolysis time according to the determined rates above. Since the binding is on the order of $2 * 10^4$ steps in the simulation, we set this to be the timestep at which hydrolysis can happen in monomers. We start the simulation with a pentamer, and keep the monomer concentration constant while introducing hydrolysis. We change the type of particles (1,2,6,7) using the functionality of fix/bond-react in LAMMPS, and then set the new $\varepsilon_H$ to the value determined from the fragmentation analysis. We then vary the hydrolysis probability $p_H$, e.i. the probability that each individual monomer inside the polymer can get hydrolysed and find we get sustained treadmilling for $p_H = 10^{-6}$.

#### 4 Implementation of the genetic algorithm

The genetic algorithm was implemented in the DEAP python package, where the internal functions - initialization, mutation and crossover were manually scripted, but the rest of the code was run using prebuilt DEAP architecture for genetic algorithms. To evolve the designs, an elitist genetic algorithm was used with parameters listed in Table 2 below.

| Parameters of the elitist genetic algorithm | | |
|---|---|---|
| Parameter | Value | Explanation |
| CXPB | 0.9 | Probability of crossover between two individuals |
| MUTPB | 0.1 | Probability of mutation of an individual |
| ngen | 50 | Number of generations in the GA |
| migrate_interval | 5 | Number of generations after which migration between demes occurs |
| num_demes | 10 | Number of demes |
| pop_size | 50 | Number of individuals in one deme |
| crosspb | 0.5 | Crossover probability of bonds in the lattice for individuals chosen for crossover |
| indpb | 0.5 | Mutation probability of bonds in the lattice for an individual chosen for mutation |
| tournsize | 5 | Number of individuals chosen for tournament selection in each deme |
| num_migrate | 5 | Number of individuals migrated every migrate_interval number of generations |
| ELITE_SIZE | 1 | Number of elite individuals in each deme |

Table 2: List of the parameters used in the genetic algorithm

The genetic algorithm developed was an elitist multi-deme genetic algorithm. Migration was performed by replacing a given number of worst performing individuals in one deme with the best performing individuals in the neighbouring deme (num_migrate from Table 2). Tournament selection was used as a selection strategy. Evaluation, crossover and mutation were performed using custom made functions to treat the individuals appropriately. There are two probabilities corresponding to mutation and crossover. One corresponds to the probability of choosing individual(s) for mutation/crossover, while the other corresponds to the probability of performing mutation/crossover on a selected first neighbour bond in the 4x4 hexagonal lattice.

The GA acts only on bonds of the monomeric structure, and it deletes particles from the monomer if there is no bond connecting it to the rest of the monomer. There are other helper functions making sure the evolved design is connected and read correctly by the LAMMPS simulation.

After evolving the monomer structure, the binding sites are chosen in the initial generation with 0.5 probability of being the top and bottom most particles and 0.5 probability of being the left and right most particles in the monomer. This is done via sampling a random number between 0 and 1 which decides between these two options for binding sites. After establishing binding sites in the initial generation, the next generation individuals inherit this random number imposing the binding site.

To evaluate the loss function, LAMMPS code was used to make a molecular dynamics simulation of monomer self-assembly. The interaction between binding sites is cosine squared with a repulsive WCA tail, with parameters shown in the table below (eps, coff). To cut down on simulation time where monomers are just diffusing, we use a narrow simulations box that follows polymer growth and we add monomers one by one to avoid growing multiple polymers at once.

Since we are interested in single polymer dynamics, we also introduce deletion in the code by removing any other polymers that might start forming in the box, or multiple monomers which can

| Parameters of the LAMMPS simulations | | |
|---|---|---|
| Parameter | Value | Explanation |
| eps | 30 kT | Attraction between binding sites |
| coff | $1.2\sigma$ | Interaction cutoff for monomer interactions |
| cutoff | $1.1\sigma$ | Cutoff used to identify monomer insertion into polymer in cluster identifying code |
| dt | 0.01 | Timestep of the Langevin integrator |
| init_region_L_height | 15 | Initial height of the simulation box |
| init_region_L_width | 15 | Initial width of the simulation box |
| delta_L | 5 | Increase in box size left/right of the seed during new monomer insertion |
| num_seeds | 10 | Number of replica simulations used to characterise the loss function of a monomer design |
| sim_steps | 2000000 | Number of simulation steps used in the GA |

Table 3: List of the parameters used in the genetic algorithm

occur through detachment from the original polymer. This also makes sure we are optimizing for kinetic polerity of the single filament, not polarity induced by blocking of one side of the filament via a different monomer or polymer. The LAMMPS code was ran through python interface, which keeps track of the clusters the monomers are forming to dynamically add new monomers and change box size. The simulation running was parallelised and tracked with the signac package from python. The simulation is run as follows:

1. The simulation starts with two monomers in a box of size init_region_L_height x init_region_L_width from Table 3
2. After the two monomers bind to form a nucleation dimer, they are fixed at the center of the box (0,0,0) via a strong spring, making sure there is no translation of the nucleation dimer. The dimer can still rotate freely in the box
3. After fixing the dimer in the center of the box, the simulation begins to track the number of monomers in solution left and right of the nucleation dimer, and keeps only one monomer on each side
4. Along with monomer addition during growth, the simulation box is increased on the side which experiences growth by delta_L from Table 3
5. An extra layer of control exists during growth, where the code detects if there is more than one polymer forming, and if so keeps only the original one and deletes the other one from the box. Similarly, if more than one free monomer exist on each side of the growing polymer, they are deleted until there is only one on each side. Each time there is a deletion event the box is shrunk to the polymer size $\pm$ 10 to avoid growing the box too much and spending long simulation times on diffusion.

The parameters used for the LAMMPS simulations are shown in the Table 3 below:
The following loss function was used to rank individuals:

$$L = \Lambda_1 N_{branches} + \Lambda_2 \exp(-\Delta COM/\Delta r)$$

where $N_{branches}$ corresponds to branching out events - calculated as the number of monomers bound to more than 2 neighbours in the simulation box. $\Delta COM$ is the shift of the center of mass (COM) of the polymer from a set monomer (monomer number 1 was used). To make sure that the structural polarity of the monomers is coupled to the kinetic polarity of the polymer, monomers were assigned an internal vector pointing from one binding site to the other. After the self-assembly was finished, the direction of the vector of the COM of the polymer was compared with the directionalities of monomers in the polymer using a cosine product. Since the LAMMPS self-assembly is run for multiple seeds and the monomers can rotate freely in the box, this makes sure that the COM shift of the polymer is consistent with the internal vector of monomers when the mean COM is taken from replicas. This coupling is set by the functional form of an exponential which is not symmetric with respect to the input, optimising for faster assembly in the internal direction of the monomer. $\Lambda_1$ and $\Lambda_2$ are weights assigned to each term in the loss, and the values used were $\Lambda_1 = 100$, $\Lambda_2 = 1000$. $\Delta r$ is a value set such that the fitness gradient is more stark and fitness values clearly indicate well performing individuals. $\Delta r = 10$ was used in the performed simulations.

To confirm kinetic polarity found in the GA, good individuals were tested in postanalysis, with the same LAMMPS self-assembly setup, but with longer simulations (sim_steps = 20000000) and larger number of simulations replicas (num_seeds = 300). To quantify kinetic polarity we track both the shift of COM as explained in the loss function calculation, as well as speed of binding on both polymer ends.

Additional analysis was performed on the evolved design depicted in Fig 2 to show that the binding area reduces as the free monomer binds to the polymer. A numerical algorithm was used to estimate the accessible binding surface of monomers. The procedure consists moving in a circle of radius equal to half of the interaction cutoff of the binding site particles and checking if there are other particles in this position (not only belonging to the monomer but any other particle in the simulation box). If there are, this means that there is a particle blocking a binding site, otherwise it is an accessible binding surface. From the information on the angles which are accessible for binding one can form a circle arc with the radius equal to the interaction cutoff. This was done in a resolution of $1^\circ$ and an example of accessible surface circilar arcs shaded in different colors representing the two binding sites for a free monomer at one timepoint are shown in Supplementary Fig 15. This procedure is then repeated for all monomers during a simulation. This allows us to gather statistics on the available binding area depending on whether the monomer is free in solution, partially bound (bound on the polymer end) or fully incorporated into the filament.

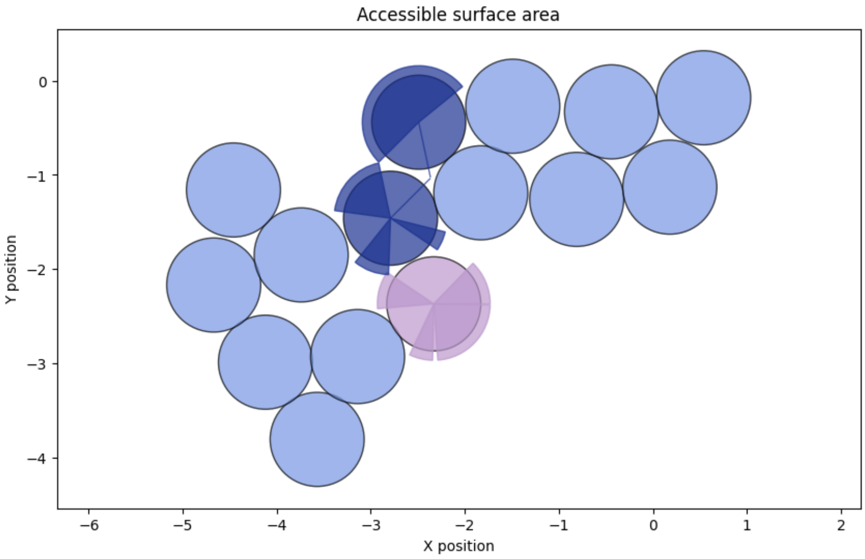

Supplementary Figure 15: Example of accessible surface calculation: the two binding sites are shown in pink and dark blue, while the shaded regions show the accessible arcs for binding to these particles

### 5 Supplementary Movies

- Supplementary Movie 1: Representative simulation showing a design evolved by the GA self-assemble into directional filaments. The monomer uses a tail-blocking mechanism to achieve kinetic polarity. Simulation parameters are: $\varepsilon = 30k_BT$. The constant concentration is kept as detailed in section 3.3, with $\Delta L = 25\sigma$, $\Delta = 15\sigma$
- Supplementary Movie 2: Representative simulation showing a design evolved by the GA self-assemble into directional filaments. The monomer uses a self-capping mechanism to achieve kinetic polarity. Simulation parameters are: $\varepsilon = 30k_BT$. The constant concentration is kept as detailed in section 3.3, with $\Delta L = 25\sigma$, $\Delta = 15\sigma$
- Supplementary Movie 3: A minimal monomer undergoing a conformation change, starting from the solution $S$-form and switching to the polymer $P$-form. Simulation parameters are: $\varepsilon_S = 6k_BT$, $\varepsilon_P = 3k_BT$.
- Supplementary Movie 4: Representative simulation showing the minimal monomers exhibiting directional growth.
- Supplementary Movie 5: Representative simulation showing a self-assembled treadmilling filament.
- Supplementary Movie 6: Representative simulation showing the monomers in the colloidal description exhibiting directional growth. Simulation parameters are: Symmetric polymerisation $\varepsilon_S = 3k_BT$, $\varepsilon_P = 6k_BT$, asymmetric polymerisation $\varepsilon_S = 6k_BT$, $\varepsilon_P = 3k_BT$, $\varepsilon = 15k_BT$. The procedure of increasing the box size is explained in detail in subsection 3.3
- Supplementary Movie 7: Representative simulation showing the monomers in the colloidal description exhibiting treadmilling. Simulation parameters are: $\varepsilon_S = 6k_BT$, $\varepsilon_P = 3k_BT$, $\varepsilon = 15k_BT$, $\varepsilon_H = 8k_BT$, $\varepsilon = 15k_BT$. Constant monomer concentration is kept in a circle around end monomers of radius $R = 8\sigma$.
